# SMART: A Spatio-Molecular Atlas of Response Trajectories in Triple-Negative Breast Cancer

**DOI:** 10.1101/2025.08.27.672673

**Authors:** Isobelle Wall, Anthony Baptista, Jelmar Quist, Holly Rafique, Mengyuan Li, Lucy Ryan, Gregory Verghese, Stefania Marcotti, Thomas A. Phillips, Claudia Owczarek, Signe Clausen, Trine Tramm, Graham Booker, Anca Mera, Jasmine Timbres, King’s Health Partners Biobank, Elinor Sawyer, Cheryl Gillet, Sheeba Irshad, Sarah Pinder, Maddy Parsons, Anita Grigoriadis

## Abstract

A major challenge in treating Triple-Negative Breast Cancer (TNBC) lies in its molecular, morphological and clinical heterogeneity, which hampers accurate prediction of responses to neoadjuvant treatment. To address this, we introduce SMART: Spatio-Molecular Atlas of Response Trajectories, a comprehensive, multimodal resource compiled from 129 TNBC samples across 89 patients, obtained before, during, and after neoadjuvant chemotherapy (NACT). SMART comprises of 5,096 high quality manually selected spatial transcriptomic profiles enriched for epithelial, immune, or stromal compartments; paralleled with histological annotations, imagebased network analysis and protein expression. Seven novel spatial epithelial archetypes (EAs), seven tumour-immune microenvironments (TIMEs) and their co-localisation patterns were defined, revealing an opposing prevalence of functionally divergent EAs between response groups and the prognostic significance of B-cell enriched TIMEs, in particular those surrounding histologically normal epithelium adjacent to the tumour. The SMART dataset and analytical tools are publicly available via the PharosAI platform, providing the research community with the most comprehensive, manually annotated spatio-molecular transcriptomics atlas of NACT-treated TNBC to date.

## 1 Introduction

Triple-Negative Breast Cancer (TNBC) is an aggressive subtype, accounting for 10−15% of all breast cancer (BC) diagnoses with the lowest five-year survival rate [1]. Neoadjuvant chemotherapy (NACT) has been the preferred approach to treat stage II or III TNBC to reduce the risk of distant recurrence and death [2]. Whilst the addition of anti-PDL1 immune-checkpoint inhibitors, namely pembrolizumab, have shown to improve overall survival (OS) in patients with early TNBC [3] the clinical benefit for patients with advanced disease is limited[4]. Moreover, as combination therapy with immune-checkpoint blockades (ICBs) and NACT become the gold standard treatment for many patients, detailed understanding of responses to NACT alone and how chemotherapy may influence the tumour-immune landscape is needed.

The intrinsic morphological and molecular heterogeneity of TNBC is thought to contribute to the variable responses to treatment, and has resulted in extensive characterisation of clinically relevant signatures[5, 6]. Large scale single-cell RNA sequencing (scRNA-seq) has revealed ‘cell-states’ which capture the dynamic physiological trajectories of distinct tumour cell phenotypes[7, 8]. The advent of spatial-omics has recently facilitated the characterisation of ‘spatial archetypes’ that capture the relative composition of cell states within spatially defined regions [9, 10]. In BC, this approach highlights the complex dynamics of tumour cell-states and the Tumour Micronenvironments (TME) across BC subtypes [5, 11] and molecular subtypes of TNBC [9, 12], with prognostic implications [9, 13]. However a spatial transcriptomics (ST) dataset guided by a priori knowledge of discrete histological morphologies alongside a scalable method to map the heterogeneity of spatial archetypes in NACT-treated TNBC is lacking.

Here, we present a Spatio-Molecular Atlas of Response Trajectories (SMART) in TNBC; a compendium of histomorphological and molecular datasets obtained from 129 TNBC samples across 89 patients before, during, and after NACT. SMART compiles rich clinical metadata, digitised whole slide H&E images with detailed histological annotations, ST profiling constituting 5,096 tumour, immune or stromal-enriched regions, immunofluorescence (IF) image-based analysis of tumour networks and imaging-mass cytometry (IMC), providing an unprecedented resource for the BC research community.

Here we position our ST dataset amongst the plethora of molecular profiling at the patient level (pseudobulk) and explore the intratumoural heterogeneity at the spatial level to identify spatio-molecular determinant of NACT response and map longitudinal changes post-NACT. We demonstrate the unique capacity of SMART to facilitate multi-modal characterisation of spatial epithelial archetypes (EAs), Tumour-Immune Micronenvironments (TIMEs) and their co-localisation. This reveals an opposing prevalence of functionally divergent EAs between response groups and the prognostic significance of B-cell enriched TIMEs. Critically, multi-modal analysis with SMART provides novel insights into the dynamics between tissue architecture and molecular profiles that correlate with distinct clinical outcomes.

## 2 Results

### 2.1 Generating A SMART Dataset

To build a SMART dataset we obtained 141 Formalin-Fixed Paraffin-Embedded (FFPE) and Fresh-Frozen (FF) tissue blocks collected from 96 patients with TNBC before during and after NACT treatment from King’s Health Partners (KHP) biobank (Guy’s Hospital, London). Overall, 27 patients received epirubicin, cyclophosphamide, and either docetaxel or paclitaxel (EC-T), 46 received EC-T in combination with carboplatin (EC-T + Carbo) and 23 were treated with alternative chemotherapy regimens (Supplementary Table 1). Patients were classified as ‘responders’ if they achieved a pathological Complete Response (pCR) or had a Residual Cancer Burden (RCB) of 0 or 1 after NACT. Conversely, ‘non-responders’ were defined as those with an RCB of 2 or higher. Out of 96 TNBC patients, 51 were classified as responders (R), 43 as non-responders (NR) and 2 received adjuvant chemotherapy (ACT). Amongst clinico-pathological features, age at diagnosis showed the strongest association with treatment response, with younger patients demonstrating higher response rates and increased statistical significance (Supplementary Table S1). Furthermore, patients under the age of 50 at diagnoses, those with ≥30% stromal Tumour Infiltrating Lymphocytes (sTILs) and histological grade 3 tumour at diagnosis demonstrated improved overall survival (Fig. S1). Additional patient characteristics are details in Supplementary Table 1.

To perform multimodal profiling of the SMART cohort, serial sections were subjected to three modalities; H&E staining, ST profiling with the GeoMX Whole Transcriptomic Atlas (GeoMX Hu WTA^TM^) and IMC. Briefly, whole slide images (WSIs) of H&E-stained sections were annotated by a consultant pathologist (SEP) to characterise the morphologies of normal epithelium (distant or tumour-adjacent), tumour (cancerisation, invasive tumour, ductal carcinoma in situ (DCIS), lymphovascular invasion (LVI), squamous epithelium and hyperplasia) and the TME (necrosis, fibrosis, stroma and inflammation), in addition to sTILs (Fig. 1). These expert annotations guided the manual selection of regions of interest (ROIs) for ST and IMC. For ST data generation, we performed uniform tiling of 250 × 250 *µ*m square ROIs across whole pre-NACT biopsies, where possible (Fig. S2 - top panel, Material & Methods). From post-NACT samples of non-responders we selected pathologist-guided 660 *µ*m-diameter ROIs from the tumour edge, tumour centre, and other histologically distinct regions e.g DCIS (Fig. S2 - bottom panel, Material & Methods). IF staining was then used to further segment each ROI into areas of illumination (AOIs) defined as PanCK+ (epithelial/tumour), CD45+ (immune), or PanCK–CD45– (stromal) compartments. Overall, we collected a total of 5,765 AOIs, with up to 172 AOIs per pre-NACT sample and up to 71 AOIs per post-NACT sample (Fig. 1, Material & Methods).

**Fig. 1:**
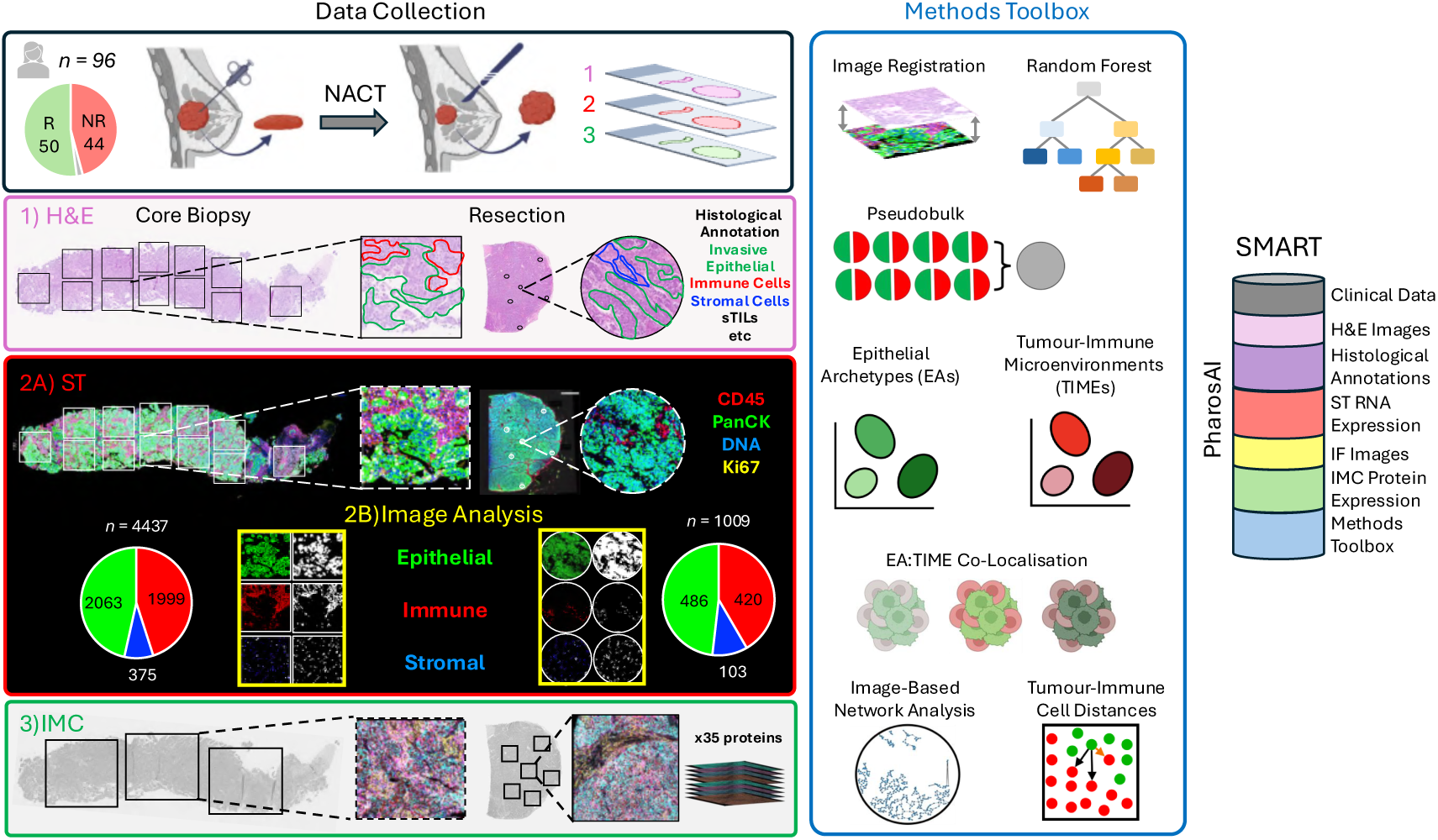
Spatio-Molecular Atlas of Response Trajectories (SMART): Multimodal, longitudinal profiling of triple-negative breast cancer (TNBC). Tissue biopsies were obtained before, during, and after neoadjuvant chemotherapy (NACT) from responder (R) and non-responder (NR) TNBC patients (black frame). Three serial sections were taken from each sample and subjected to distinct analytical modalities. Section 1 was stained with H&E (pink frame), imaged, and annotated by a pathologist to identify histological features and quantify stromal tumour-infiltrating lymphocytes (sTILs). Section 2 underwent spatial transcriptomics (ST) profiling (red frame) using the NanoString/Bruker GeoMx Digital Spatial Profiler (DSP) to capture RNA transcripts from epithelial (PanCK+) and immune (CD45+) cells for downstream analysis. Immunofluorescent (IF) images for each AOI were obtained from the GeoMx DSP (yellow frame) and subjected to downstream imagebased network analysis. Section 3 was analysed using imaging mass cytometry (IMC) (green frame) to quantify the expression of 35 protein markers. The methods toolbox (blue frame) displays all analytical tools developed in this study. The SMART dataset compiles all datasets and analytical methods which are hosted on PharosAI and made publicly available (See Data Availability).

After a rigorous filtering with our custom quality control (QC) pipeline (Fig. S2, Material & Methods), we retained high-quality ST data from 5,096 AOIs with an overall drop-out rate of 11.6% (Fig. S3) across 129/141 tumours from 86/89 patients (R = 46, NR = 46, ACT = 2; Fig. S3). From each AOI, RNA expression of more than 18,000 genes was quantified and analysed alongside histological annotations, IF image-based network models and cell-cell distance calculations, as well as IMC expression data of 35 proteins from parallel ROIs (Fig. 1, Material & Methods). The SMART dataset is publicly available through PharosAI platform, and the tools on the following Github repository: github/izziWall/SMART.

### 2.2 SMART Recapitulates Established Prognostic Signatures In TNBC

Bulk transcriptomics studies have previously sought to identify transcriptional profiles of distinct TNBC subtypes with prognostic significance. To date, several gene-expression based classifiers have been established, namely the PAM50 classifier of intrinsic breast cancer subtypes [14], the Lehmann classifier [6] and the Baylor classifier[15]. To investigate the distribution of TNBC subtypes in our cohort, we first used the SMART dataset to identify and extract PanCK+ AOIs from histomorphologically ‘invasive’ regions (Material & Methods). These regions were subsequently aggregated across individual tumours (Fig. S4, Material & Methods) to generated a pseudobulk dataset of invasive TNBCs. Of 129 TNBC cases, 70 pre-NACT (R = 42, NR = 28) and 39 post-NACT (R = 0, NR = 39) pseudobulk samples passed QC and were subjected to molecular classification (Material & Methods).

The PAM50 classifier identified the majority as basal-like (R = 35/42, NR = 21/28), followed by the HER2-enriched subtype (R = 6/42,NR = 6/28; Fig. 2A, left panel). Notably, the Lehmann classifier found the basal-like-1 (BL1) subtype to be enriched in responders (R = 11/42) compared to non-responders (NR = 3/28). By contrast, the basal-like-2 (BL2) subtype was more prevalent in non-responders (NR = 4/28) compared to responders (R = 1/42; Fig. 2A - middle panel), consistent with previous reports [16]. The Baylor classifier showed a slight enrichment of the basal-like immune-activated (BLIA) subtype in responders (R = 21/42) compared to non-responder (NR = 11/28), meanwhile the basal-like-immune-suppressed (BLIS) subtype was comparable between response groups (R = 18/42, NR = 13/28; Fig 2A, right panel).

**Fig. 2:**
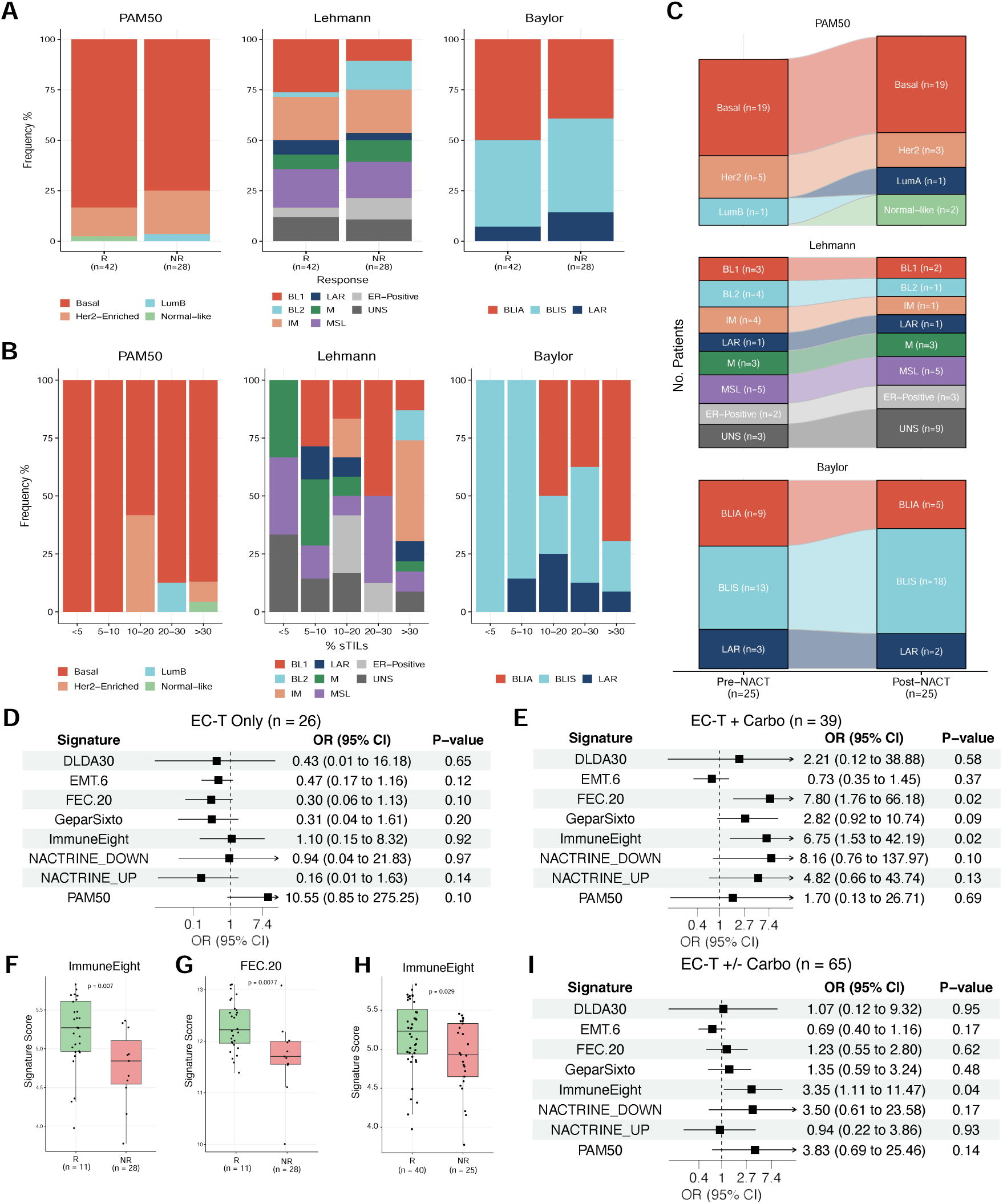
Pseudobulk Analysis Of Spatial Transcriptomics Data Recapitulates Prognostic Features In TNBC: Stacked bar charts illustrate the proportion of TNBC subtypes according to the PAM50 (left panel), Lehmann (middle panel), and Baylor (right panel) classifiers, in pre-NACT samples stratified by **A)** response group and **B)** percentage of stromal TILs. **C)** Alluvial plots show changes in TNBC subtypes between preand post-NACT samples, according to the PAM50 (top panel), Lehmann (middle panel), and Baylor (bottom panel) classifiers. Forest plots indicating the odds ratio (OR), 5% and 95% confidence intervals, and p-values for TNBC gene signatures associated with response to NACT in a cohort of patients treated with **D)** ECT only, **E)** ECT + carboplatin and **I)** ECT ± carboplatin. Boxplots indicate signature scores in responders (green) and non-responders (red) that are strongly associated with response to treatment in **F & G)** ECT + carboplatin or **H)** ECT ± carboplatin. The number of patients in each response group is indicated below the x-axis. Wilcoxon test *p*-values are shown at the top of each boxplot; statistical significance was defined as *p <* 0.05.

TNBC is considered to be more immunogenic, with levels of sTILs being significant predictors of responses to both NACT and immunotherapy [17, 18]. Therefore, we next asked whether the distribution of TNBC subtypes changes with increasing infiltration of sTILs[19, 20]. Overall, sTILs scoring was performed on digitised H&E stained slides of pre-NACT biopsies from 66/96 TNBC patients (Materials & Methods), of which 39 (R = 17, NR = 21, ATC = 1) were classified as having low sTILs (*<*30%) and 27 (R = 15, NR = 11, ATC = 1) had high sTILs (≥30%) (Supplementary Table 1). The PAM50 classifier did not distinguish between patients with high versus low sTILs (Fig. 2B, left panel). As expected, the Lehmann IM and Baylor BLIA subtypes each accounted for the highest proportion of TNBCs with ≥30% (Fig. 2B) [21]. Meanwhile, TNBCs with sTILs *<*30% were largely defined by the Baylor classifier as BLIS subtypes (Fig. 2B, right panel).

Given the known influence of chemotherapy on transcriptional heterogeneity [22] and Lehmann TNBC classification[23], we next investigated how TNBC subtypes change in 25 patient-matched pre-NACT and post-NACT pairs. Overall, 36%, 56% and 20% of patients demonstrated a change in TNBC subtype according to the PAM50, Lehmann and Baylor classifiers, respectively (Fig. 2C). The PAM50 classifier found the most frequent change to occur in HER2-enriched subtypes which adopted a Basal-like phenotype following NACT (n = 4/5). Meanwhile those classified as Basal-like in the neoadjuvant setting adopted HER2-enriched (n = 2/4) and Normal-like (n = 2/4) phenotypes post-NACT (Fig. 2C - top panel). By contrast, the response trajectory of Lehmann TNBC subtypes was highly variable with no consistent pattern observed(Fig. 2C, middle panel). In accordance with the immunomodulatory effects of chemotherapy [24], the Baylor classifier found 4/9 pre-NACT biopsies with a BLIA subtype to transitioned to a BLIS phenotype (Fig. 2C - bottom panel).

### 2.3 Immune-related gene signatures differentiate responders from non-responders in a SMART pseudobulk dataset

We next sought to explore the performance of previously reported predictive signatures in the pseudobulk setting, including the PAM50 proliferation signature [25, 26], a 20-gene classifier (FEC.20)[27], an 8-gene panel (ImmuneEight) [28], the GeparSixto immune signature score [29], two signatures (NACTRINE UP, NACTRINE DOWN) prognostic for pCR in the NACTRINE trial [30], a 6-gene signature relating to epithelial-mesenchymal transition (EMT) [31] and a pharmacogenomic predictor of chemotherapy sensitivity (DLDA30) [32]. To do so, we aggregated counts from PanCK+, CD45+, and PanCK-/CD45-AOIs across individual pre-NACT biopsies to generated a ‘global’ pseudobulk dataset (Fig. S4) and calculated a sample-level score for each signature (Material & Methods).

To ensure accurate modelling of treatment response, given the known clinical benefit of adding carboplatin to standard chemotherapy on overall survival in TNBC [33], we stratified the input data into three treatment cohorts: (1) patients treated with EC-T alone (n = 26), (2) those who received EC-T in combination with carboplatin (n = 39), and (3) a combined cohort of all patients who received EC-T with or without carboplatin (EC-T ± Carbo; n = 65). We then applied a linear regression model to assess the association between each signature score and binary-coded responses to NACT (Responder = 0, Non-Responder = 1).

Out of the eight signatures assessed across the three cohorts, only two were found to be significantly associated with responses to NACT. Whilst no signature was significantly associated with outcome in the ECT-only cohort (Fig. 2D), an increased expression of both the ImmuneEight and the FEC.20 signature were associated with improved responses to treatment with ECT + Carboplatin (Fig. 2E-G). Notably, the statistical significance of the ImmuneEight signature persisted in the combined cohort of ECT ± Carboplatin however that of the FEC.20 signature was lost (Fig.2H & I).

Together, these findings demonstrate the capacity of SMART to facilitate histologically informed pseudobulk analysis of an ST dataset, demonstrate the profound influence of NACT on molecular profiles of TNBC and reaffirm the prognostic significance of immune-related signatures in response to NACT.

### 2.4 Limited ROI subsets retain predictive accuracy for treatment response in TNBC

SMART confers the unique benefit of performing both patient level and spatially-resolved analysis. Therefore, we sought to interrogate the clinical significance of spatially-resolved expression data. To explore the prognostic potential of our ST dataset, we trained a Random Forest classifier to distinguish between ROIs collected from responders and non-responders. The model (Fig. S5A) was trained on normalized expression values of the 1,000 most variable genes from tumour-associated ROIs (see Material & Methods). The Random Forest did not consistently identify a single predictive set of ROIs, suggesting that multiple, distinct ROIs have comparable predictive power (Fig. S5B). However, based on a learning curve analysis (Fig. S5C), we showed that approximately 121 bootstrapped ROIs are sufficient to achieve 75% classification accuracy, indicating that a limited yet informative subset of ROIs can effectively predict treatment response in TNBC despite its molecular heterogeneity.

### 2.5 Epithelial Archetypes (EAs) Capture Transcriptional Cancer Cell-States Related to Histological and Network Metrics

To explore the spatio - molecular features of epithelial cells in the SMART dataset, we use *Signifinder* [34] to first quantify the expression of published gene signatures relating to cancer cell states in each PanCK+ AOI which were then subjected to consensus clustering (Fig. S6). We subsequently identified seven stable clusters of PanCK+ AOIs, referred to hereafter as ‘epithelial archetypes’ (EAs), characterised by the relative expression of 26 cell state signatures relating to EMT, cell cycling, cell death, responses to immune checkpoint blockade (ICB), immune responses, chemokine signalling, chromosomal instability (CIN), extracellular matrix (ECM) remodelling, hypoxia, angiogenesis, autophagy, stemness and TGF-*β* programming of cancer-associated fibroblasts(Fig. 3A-B).

**Fig. 3:**
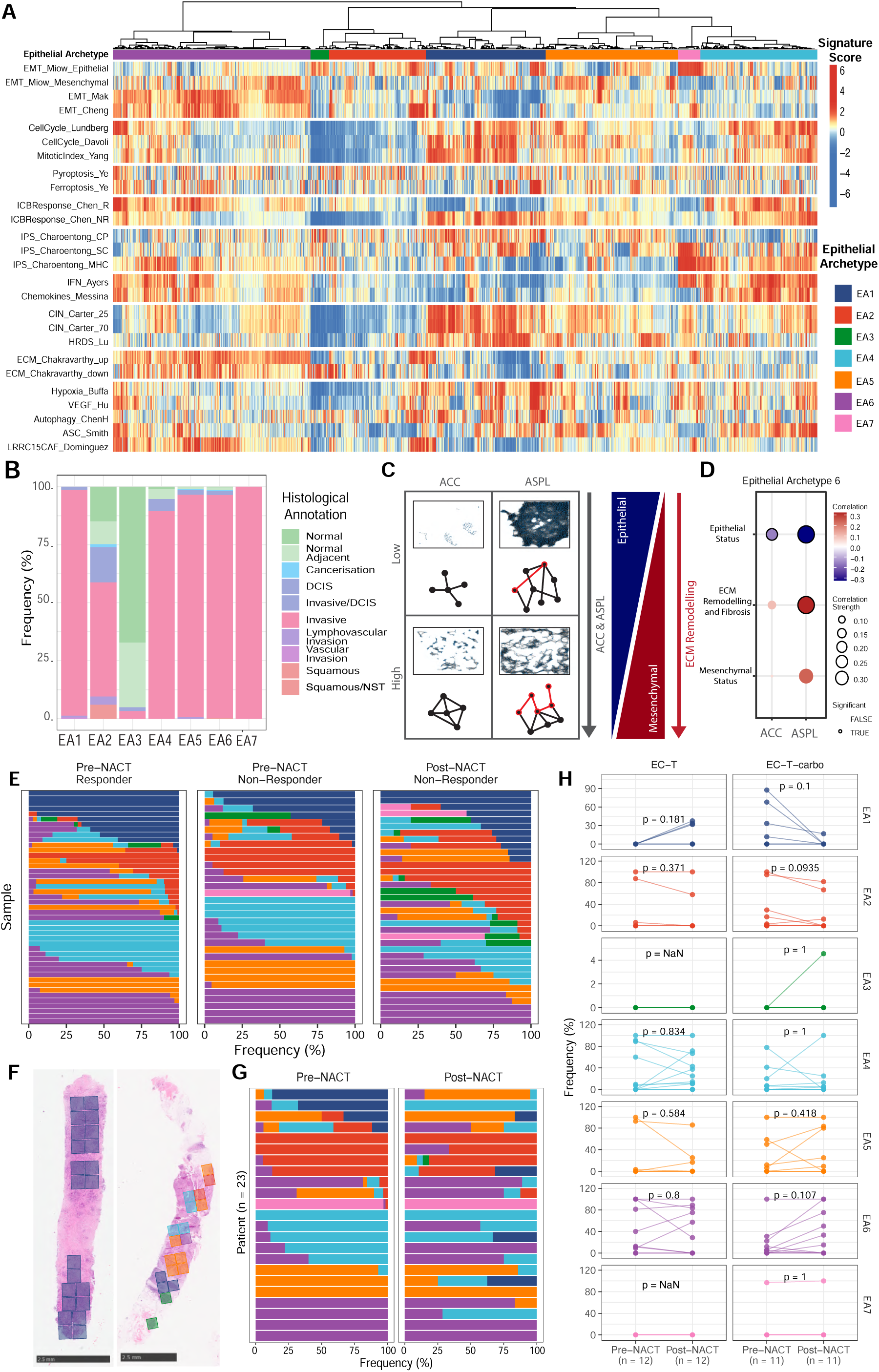
Characterisation and Distribution of Epithelial Archetypes: **A)** Heatmap depicts relative enrichment of selected signature scores per sample. Samples are clustered according to *k* = 7 as determined using consensus clustering and coloured bars indicate the assignment of each sample to an ‘epithelial archetype’ (EA), labelled below the heatmap. The relative expression is given on a colour scale ranging from blue (low expression) to red (high expression). **B)** Stacked bar chart displaying the distribution of different histological annotations associated with each sample per EA. **C)** Schematic diagrams and network graphs from EA6 AOIs with the lowest (top panel) and highest (bottom panel) average clustering coefficient (ACC) and average shortest path length (ASPL). Parallel schematics indicate that as each of ACC and ASPL increase, the expression of signatures related to the epithelial-status of EA6 decreases whilst that of the mesenchymal-status and ECM remodelling increases. **D)** Dotplot indicating the Pearson correlation between network metrics and EMT-related signatures in ROIs from EA6. Statistical significance was considered when *p <* 0.05. **E)** Horizontal stacked bar charts showing the frequency of EA per pre-NACT sample from responders (n = 45, left panel), non-responders (n = 31, middle panel) and post-NACT samples from non-responders (n = 38, right panel). The colours of each bar segment indicate the corresponding EA as represented in panel A. **F)** Representative H&E images of pre-NACT biopsies overlaid with GeoMx ROIs coloured according to their assigned EA. **G)** Horizontal stacked bar charts indicating changes in EA frequencies in patient-matched pre-NACT and post-NACT samples, where each row indicates one patient. **H)** Patient-matched dot plots indicating the change in EA frequencies from 23 pre-NACT and post-NACT pairs. Paired wilcoxon test *p*-values are given above the plot, significance was defined as *p <* 0.05.

EA1 was primarily characterised by high enrichment of cell cycle and CIN signatures in addition to hypoxia, VEGF and autophagy signatures (Fig. 3A). By contrast, EA2 and EA3 were characterised by a downregulation of signatures related to cell cycle, CIN, hypoxia and VEGF. EA4 was characterised by immune-related signatures including antigen processing and presentation by MHC molecules, IFN and chemokine signalling. EA5 was characterised by a moderate expression of cell cycle and CIN related signatures, relative to EA1 and EA2. EA6 was uniquely characterised by the upregulation of EMT and ECM-related signatures, often accompanied by IFN and chemokine signalling (Fig. 3A). EA7 was enriched for signatures relating to cellular senescence, antigen processing and presentation by MHC molecules, IFN signalling and an epithelial-like status with regards to EMT (Fig. 3A). We then asked what were the histomorphological features of each EA and found that EA1 and EA4-7 almost exclusively comprised of ‘invasive’ tumour cells (Fig. 3B). Interestingly, EA2 demonstrated a unique enrichment for regions of squamous epithelia or lymphovascular invasion, whilst also capturing regions of normal/normal adjacent epithelia, cancerisation and invasive tumour (Fig. 3B). Meanwhile, EA3 was comprised almost exclusively of normal and normal-adjacent epithelia (Fig. 3B). Given that EA6 was characterised by EMT enrichment and ECM-remodelling, which are associated with mechanical and structural changes in invasive breast cancer [35], we asked if the transcriptional profile of EA6 correlated with topological changes in the tissue. To this end, we conducted image analysis of PanCK+ AOIs from EA6 to calculate two graphical network metrics that capture geometric patterns within the tissue. The average clustering coefficient (ACC), to measure the degree of clustering amongst tumour cell neighbourhoods, and the average shortest path length (ASPL) to measure the number of ‘steps’ between each tumour network (Fig. 3C, Methods & Materials); representing the connectivity and dissemination of tumour networks, respectively. For example, an ROI with a high ACC and high ASPL would indicate a network of tightly formed tumour cell clusters that are far apart from each other (Fig. 3C). We next computed the Pearson correlation between each network metric and the expression of signatures relating to the epithelial status, ECM-remodelling and mesenchymal status of EA6. Both the ACC and ASPL negatively correlated with the epithelial-like status of tumour cells, meanwhile the ASPL was positively correlated with ECM remodelling and the mesenchymal status of tumour cells (Fig. 3D).

We next evaluated the distribution of EAs across individual tumour samples and found that no EA consistently occurred in all samples and some were more heterogeneous than others (Fig. 3E-F). In pre-NACT biopsies, we observed an increased prevalence of EA1 in responders compared to non-responders, meanwhile the opposite trend was observed for EA2 (Fig. 3E). Paired analysis of patient-matched samples revealed that whilst EA1 was frequently depleted post-NACT (Fig. 3G), this was exclusive to patients that received ECT + Carboplatin (Fig. 3H), meanwhile EA1 was increased in those treated with EC-T alone (Fig. 3H).

### 2.6 Deconvolution of immune cell AOIs adjacent to epithelial cells identifies subsets associated with NACT response

To characterise the immune landscape of each ROI, we applied the *SpatialDecon*[36] method to estimate immune cell abundances in CD45+ AOIs, which were then subjected to consensus clustering (Fig. S7). We subsequently identified seven stable CD45+ clusters, referred to hereafter as ‘Tumour-Immune Microenvironments’ (TIMEs) characterised by differential proportions of immune cell subtypes(Fig. 4A). Three TIMEs were uniquely dominated by memory CD4+ T cells (TIME1), plasma cells (TIME2), and naive CD8+ T cells (TIME6). By contrast, TIME3 was characteristic of a composite B cell niche, comprising of naive B cells, memory B cells, and plasma cells co-localised with plasmacytoid dendritic cells (pDCs). TIME4 was highly enriched for memory CD8+ T cells and myeloid dendritic cells (mDCs). TIME5 was almost exclusively comprised of myeloid cells, including macrophages, mast cells, NK cells, monocytes, and neutrophils, while exhibiting almost no lymphoid cells. Finally, TIME7 represented a cluster enriched for macrophages, naive CD4+ T cells, monocytes, and endothelial cells. Serial sections analysed by IMC and digitised H&E images further validated the presence of distinct cell types in TIMEs 1-5 (Fig. S8).

**Fig. 4:**
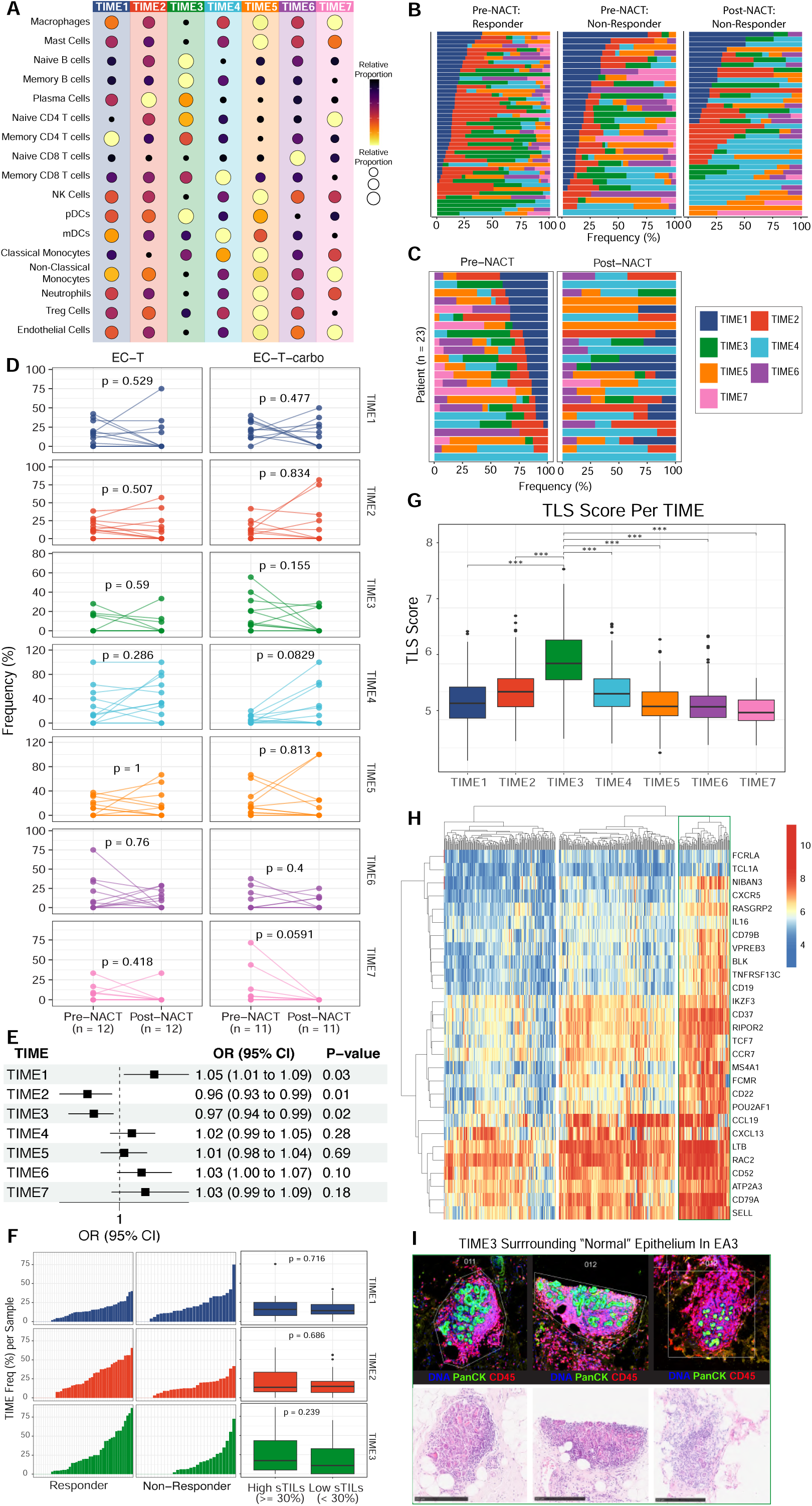
Tumour-Immune Microenvironments (TIMEs) Enriched For B cells and CD4 T cells Are Prognostic For Responses To NACT: **A)** Dot plot indicating the relative proportion of immune cell subsets per TIME cluster, determined by consensus clustering with *k* = 7. **B)** Horizontal stacked bar charts showing the frequency of TIME states per pre-NACT sample from responders (n = 45, left panel), non-responders (n = 30, middle panel) and post-NACT samples from nonresponders (n = 36, right panel). **C)** Horizontal stacked bar charts indicating changes in TIME distribution in patient-matched pre-NACT and post-NACT samples, where each row indicates one patient. **D)** Patient-matched dot plots indicating the change in TIMEs frequencies from pre-NACT to post-NACT samples. Paired wilcoxon test *p*-values are given above the plot, significance was considered as *p <* 0.05. **E)** Forest plot showing the odds ratio (OR), 5% and 95% confidence intervals and *p*-values for TIME frequencies associated with responses to NACT. **F)** Barcharts depicting the relative frequencies of TIME1 (top panel), TIME2 (middle panel) and TIME3 (bottom panel) in pre-NACT samples between responders and non-responders (left panel) and between patients with high sTILs (*>* 30%) versus low sTILs (*<* 30%)(right panel). Wilcoxon test *p*-values are provided within the plot, significance was defined as *p <* 0.05. **G)** Boxplot indicating the TLS score of CD45+ AOIs in each TIME. Pairwise comparisons between TIMEs were performed with p-values adjusted for multiple testing using the Benjamini-Hochberg (BH) method. Statistical significance is denoted as follows: *p <* 0.001 (***), *p <* 0.01 (**), *p <* 0.05 (*). Asterisks correspond to adjusted *p*−values. **H)** Heatmap displaying the relative expression of each gene from the TLS signature (reference is in the overleaf version) in CD45+ AOIs in TIME3, cut to three trees. The relative expression is given on a colour scale ranging from blue (low expression) to red (high expression). The green box indicates a cluster of TIME3 AOIs with high expression of nearly all genes in the TLS signature. **I)** Representative GeoMx IF (top panel) and matched H&E (bottom panel) images corresponding to TIME3 AOIs identified in panel G (far-right tree, green box). GeoMx IF images visualise DNA (blue), PanCK+ cells (green) and CD45+ cells (red) identifying the nuclei, epithelial cells and immune cells, respectively.

Notably, we observed increased frequencies of TIME1 in pre-NACT samples of non-responders compared to responders, meanwhile TIME2 and TIME3 were enriched in responders (Fig. 4B). Following NACT, we observed an overall increase in TIME2 and TIME4 (Fig. 4B), however paired analysis revealed a more heterogeneous response. Whilst TIME1 was markedly downregulated post-NACT in some patients, others demonstrated no change or an increase in TIME1 (Fig. 4C-D). Notably, TIME4 was shown to increase consistently in response to treatment with EC-T plus carboplatin (Fig. 4D). Analysing one example of patient-matched biopsies obtained before and during sequential cycles of NACT revealed a marked depletion in epithelial cells in parallel with contrasting patterns in the immune compartment (Fig. S9 A & B). Specifically, CD8+ T cells were markedly increased after just one cycle of NACT, followed by an a slight decrease after a second cycle (Fig. S9B). CD4+ T cells however showed a sequential increase with each cycle while memory B cells were increased only after two full cycles (Fig. S9 B). These results are consistent with ‘on-treatment’ and post-NACT analyses of TIL phenotypes in breast cancer [24] which reflect the immunogenic properties of chemotherapy. Meanwhile all other subtypes were either unchanged or depleted after two NACT cycles (Fig. S9B).

Considering the observed differences in TIME distributions between response groups, we fitted a linear regression model to assess the association between TIME frequencies per pre-NACT sample and responses to NACT. This revealed a significant association of TIME2 (plasma cells) and TIME3 (B cells) with a favourable response to NACT (Fig. 4E). In contrast, higher frequencies of TIME1 (CD4 Memory T cells) was associated with a poor response to NACT (Fig. 4E).

sTILs are known predictors of pCR following NACT [37], therefore, we asked whether the prognostic significance of TIME1, TIME2, or TIME3 was driven by different levels of sTILs, however no significant differences were observed in their relative frequencies between patients with overall high (≥ 30%) versus low (*<* 30%) sTILs (Fig. 4F). The association of TIME3 with responses to NACT is concurrent with the prognostic significance of B cells [38] and tertiary lymphoid structures (TLS) [39] in breast cancer. TLS are organised immune cell aggregates that mirror the structure and function of germinal centres (GC) in non-lymphoid tissue, exhibiting a prototypical B cell zone surrounded by a T cell zone and a collar of antibody-producing plasma cells [40]. Given that TIME3 was characterised by an enrichment for B cells at varying stages of differentiation with moderate CD4+ T cell enrichment, we hypothesised that TIME3 could be indicative of TLS. We therefore utilised a TLS signature derived from a spatial transcriptomics TNBC atlas [9] to further investigate this hypothesis. As expected, TIME3 demonstrated a significantly higher TLS score compared to all other TIMEs (Fig. 4G), however a sub-cluster of TIME3 AOIs were identified with consistently high expression of all genes in the TLS signature (Fig. 4H). Inspecting the IF and H&E images of the corresponding ROIs revealed that many of these TLS-like TIME3 AOIs surrounded histologically normal breast lobules (Fig. 4I), rather than isolated immune cell aggregates characteristic of TLS. This motivated us to further investigate the co-occurrences of each EA and TIME state.

### 2.7 Co-Localisation of EAs and TIMEs Reveal Distinct Spatial Patterns & Functional Differences Between Response Groups

To explore the co-occurrence of EAs and TIMEs within the same ROI, we first filtered our dataset to include only ROIs containing both PanCK+ (epithelial) and CD45+ (immune) AOIs that were assigned an EA or TIME state, respectively. We then applied permutation testing to identify statistically enriched pairwise combinations of EAs and TIMEs, stratified by response group and time point(Fig. 5A).

**Fig. 5:**
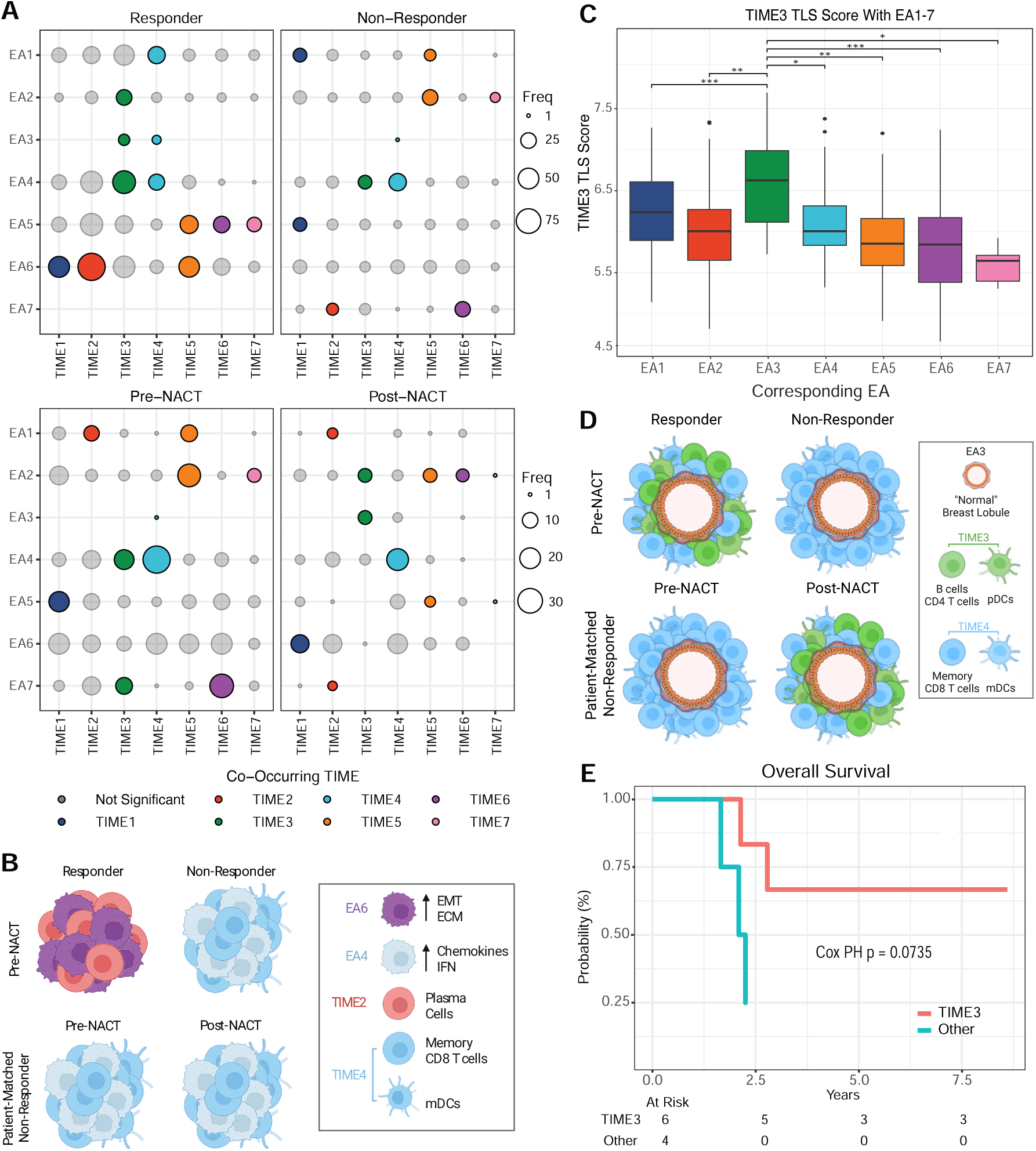
SMART Reveals The Co-Occurrence of EAs and TIMEs With Clinical Implications. **A)** Dot plots comparing statistically probable co-occurrence of EA and TIME in pre-NACT samples between responders versus non-responders, and in non-responders between pre-NACT versus post-NACT samples. The size of each dot indicates the frequency of the co-occurrence. Statistically significant co-occurrences are filled according to the corresponding TIME whilst non-significant ones are shown in grey. **B)** Schematic representation of the most frequent co-occurrences in pre-NACT samples form responders and non-responders, and in patient-matched pre-NACT and post-NACT samples from non-responders. **C)** Boxplot comparing the TLS scores of TIME3 AOIs when they co-occur with different EAs. Pairwise comparisons between TIME3 TLS scores corresponding to each EA were performed with p-values adjusted for multiple testing using the Benjamini-Hochberg (BH) method. Statistical significance is denoted as follows: p *<* 0.001 (***), p *<* 0.01 (**), p *<* 0.05 (*). Asterisks correspond to adjusted p-values. **D)** Schematic representation of EA3 co-occurrences in pre-NACT samples form responders and non-responders, and in patient-matched pre-NACT and post-NACT samples from non-responders. **E)** Kaplan-Meir survival curves comparing distant metastasis-free survival (DRFS) of non-responders when EA3 is present post-NACT and co-occurs with (red) or without (green) TIME3. Statistical significance was calculated using cox proportional hazard model.

Overall, 45/49 possible combinations of EAs and TIMEs occurred at least once, with several co-occurrence identified as statistically probable (Fig. 5A). In pre-NACT biopsies from responders, the most frequent co-occurrence was EA6 with TIME2 meanwhile EA4 with TIME4 was the most frequent co-occurrence in non-responders both before and after NACT (Fig. 5A-B). Additionally, this analysis revealed distinct co-localisation patterns of TIMEs that were previously shown to be associated with responses to NACT. In the neoadjuvant setting, significant co-localisation patterns of prognostic TIMEs were as follows: TIME1 with EA6 in responders but with EA1 and EA5 in non-responders, TIME2 with EA6 in responders but with EA7 in non-responders, and TIME3 with EA2-4 in responders but only EA4 in non-responders (Fig. 5A - top panel). However in patient-matched pairs from non-responders preand post-NACT, those patterns were as follows: TIME1 with EA5 pre-NACT but with EA6 post-NACT, TIME2 with EA7 both before and after NACT, and TIME3 with EA4 & EA7 pre-NACT but with EA2 & EA3 post-NACT (Fig. 5A - bottom panel). The co-localisation of TIME3 (enriched for B cells, CD4+ T cells and pDCs) with EA3 (comprised almost exclusively of ‘normal’ epithelium - Figure. 3B) was reminiscent of the previously observed enrichment of a TLS-like niche around normal tissues (Fig. 4H-I) and further analysis revealed that TIME3 TLS scores were significantly higher when co-localised with EA3 compared to all other EAs (Fig. 5C). This is consistent with reports of ‘immature’ or ‘early’ TLS, characterised by tight aggregates of B and T cells that lack a central GC [41], surrounding pre-malignant lesions of other cancer types [42]. Whilst TIME3 co-localised with EA3 in pre-NACT biopsies of responders, this pattern was only seen to emerge in non-responders after NACT (Fig. 5A & D). We therefore asked if the emergent co-localisation of TIME3 with EA3 in residual disease conferred a survival benefit. We found that the seven patients with at least one co-occurrence of TIME3 with EA3 had prolonged overall survival (OS) compared to the four patients EA3 existed alone or with other TIMEs (Fig. 5E), however statistical significance was not achieved, possibly due to the limited number of patients. Nevertheless, on account of the strong trend observed, we hypothesise that TIME3 captures the presence of ‘immature TLS’ around potentially pre-malignant epithelium that is prognostic for NACT responses in the neoadjuvant setting, and protective against distant recurrence in residual disease. However, additional studies are required to test this hypothesis further.

Additionally, we found that TIME3 and TIME4 consistently co-localised with EA4 in both responders and non-responders groups (Fig. 5A). Whilst TIME3 (B cells, CD4+ T cells and pDCs) was associated with response to NACT, TIME4 (Memory CD8+ T cells) was not, despite previous reports of CD8+ T cells’ prognostic significance in TNBC [43]. We therefore asked if the relative proportions of immune cell subtypes in TIME3 and TIME4 differed between response groups. When TIME3 co-localised with EA4, the relative proportion of naive CD4+ T cells and pDCs was lower in non-responders compared to responders, meanwhile that of plasma cells was increased (Fig. 6A). When TIME4 co-localised with EA4, pDCs were found to be depleted in non-responders (Fig. 6B). Since EA4 is characterised largely by the expression of several chemokines, we then asked if EA4-derived signalling could be responsible for these observed differences. Many chemokines were positively correlated with immune cell subtypes in TIME3 and TIME4 that express their cognate receptor, for example CXCL10 with memory CD8+ T cells and CXCL13 with memory B cells (Fig. 6C). However, whilst CCL19 and CCL21 are well known mediators of type 1 and type 2 T cell responses, no significant correlation was observed between their expression and the relative abundance of naive CD4+ T cells (Fig. 6C). Given that pDCs present antigens to both B cells [44] and CD8+ T cells [45], we asked if the proportion of pDCs correlated with the activation, proliferation and differentiation of B and T cells in TIME3 and TIME4. Despite a strong positive correlation between the proportion of pDCs and antigen presentation and processing in TIME3 when co-localised with EA4 in non-responders, there was a strong negative correlation with B cell proliferation (Fig. 6D, left panel). Whilst there was a moderately positive correlation between pDCs and CD8+ T cell proliferation in TIME4 when co-localised with EA4, a strong negative correlation was observed with CD8+ T cell differentiation (Fig. 6D, right panel).

**Fig. 6:**
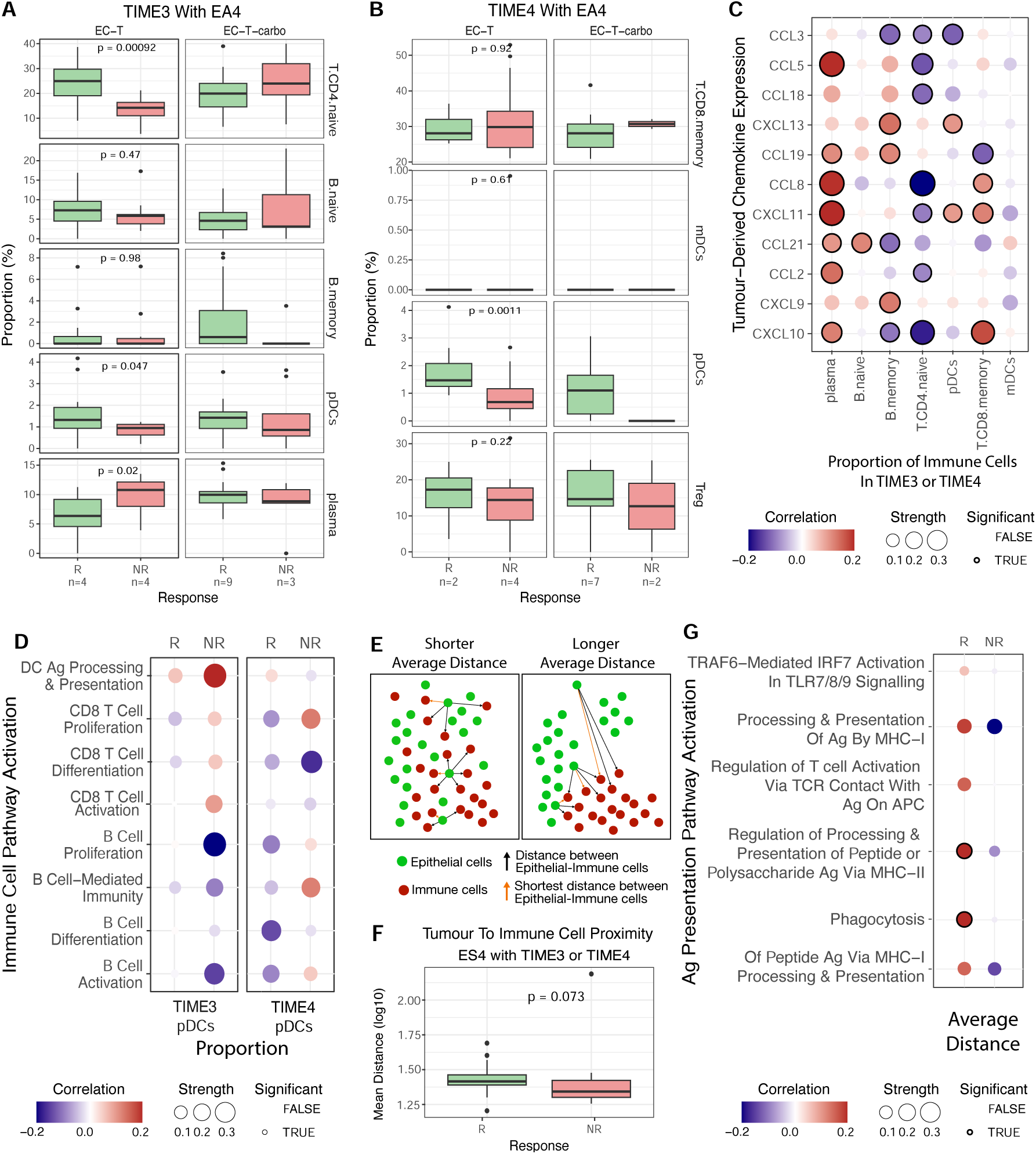
Exploring Tumour-Immune Interactions With A SMART Dataset: Boxplots indicate the relative proportion of immune cell subtypes in **A)** TIME3 AOIs co-localised with EA4 and **B)** TIME4 AOIs co-localised with EA4, between responders and non-responders. Wilcoxon test *p*-values are given above the plot, significance was defined as *p <* 0.05. The total number of patients is given below the plot. **C)** Dot plot depicts the correlation between tumour-derived chemokines expressed in EA4 with the proportions of immune cell subtypes in the corresponding TIME3 or TIME4 AOIs. Statistical significance was considered when p¡0.05 and is indicated by a thick black outline. **D)** Dotplot indicates the Pearson correlation between the relative proportions of pDCs and activation of pathways related to B-cell and CD8+ T cell stimulation within TIME3 (left panel) or TIME4 (right) AOIs when co-localised with EA4, in responders compared to non-responders. Statistical significant was considered when p *<* 0.05 and is indicated by a thick black outline. **E)** Schematic depicting the mean distance between epithelial (green) and immune (red) cells. **F)** Boxplot comparing the average distance between tumour (PanCK+) and immune (CD45+) cells in ROIs with EA4 localised with TIME4. Wilcoxon t-test was used to compute statistical significance. **G)** Dotplot indicates the correlation between the average distance between tumour (PanCK+) and immune (CD45+) cells and the relative enrichment of pathways related to antigen processing and presentation by antigen presenting cells (APCs) in TIME3 or TIME4 AOIs when localised with EA4. Statistical significant was considered when p *<* 0.05 and is indicated by a thick black outline.

We next investigated whether differences in the proximity of immune cells to tumour cells between response groups might contribute to poor responses to NACT. We therefore computed the mean distance between tumour (PanCK+) and immune (CD45+) cells within each ROI (Fig. 6E, Material & Methods). Overall, mean distances varied between each EA, with EA3 showing reduced distances in responders compared to non-responders, meanwhile EA2 and EA4 demonstrated increased distances in non-responders (Fig. S10). In ROIs comprised of EA4 and TIME3 or TIME4, specifically, the mean distance was slightly increased in responders compared to non-responders (Fig. 6F). Since the proportion of pDCs negatively correlated with B cell proliferation in TIME3 of non-responders but not responders, and T cell differentiation in TIME4, we asked if the mean distance between tumour and immune cells correlated with the function of antigen presenting cells (APCs). In responders, we found a significant positive correlation between mean distances and the enrichment of pathways related to APC function in TIME3 and TIME4 of responders (Fig. 6G), by contrast we observed a significant but negative correlation in non-responders (Fig. 6G). Together, these findings point to a highly complex TME in which activation of effector B and T cells by professional APCs may be dysregulated when TIME3 and TIME4 co-localise with EA4, however, further scrutiny of cell-cell interactions is required to elucidate these exact mechanisms. Moreover, these results indicate the unique capacity of the SMART dataset to explore tumour-immune cell interactions within defined spatio-molecular niches between response groups and following treatment with systemic chemotherapy.

## 3 Discussion

TNBC remains highly challenging due to its molecular heterogeneity, poor prognosis, and limited therapeutic options. Spatial transcriptomics (ST) has emerged as a powerful tool to interrogate TNBC heterogeneity, yet most studies to date are restricted by small patient cohorts and often lack integration of tissue architecture, morphology and proteomics. While recent large-scale profiling, such as Wang *et al* ‘s study of 94 primary tumours treated with adjuvant therapy[9], has provided important insights into clinical outcomes, critical knowledge gaps remain regarding spatio-molecular patterns that determine responses to NACT and how these are altered in residual disease. To address these gaps, we present our SMART dataset, compiling comprehensive clinical metadata, H&E-stained digitised tissue images with detailed histological annotations, ST data comprising of 5,096 manually selected AOIs, enriched for epithelial, immune or stromal cells, alongside matched image-based analysis of tissue architecture, and protein expression data for 129 TNBC tumours obtained from 89 patients before and after NACT. To the best of our knowledge, this represents the largest longitudinal pathologist-annotated ST atlas of TNBC to date. Our study demonstrates that ST can resolve the complex interplay between tumour architecture, immune infiltration, and therapeutic response in NACT TNBC. By integrating spatially resolved gene expression with histological context, we identified distinct tumour-immune niches that persist post-NACT and are enriched in non-responders, suggesting spatially confined mechanisms of chemoresistance and underscoring the prognostic and therapeutic relevance of spatial heterogeneity. These findings position ST as a powerful tool for stratifying patients beyond conventional pathological response, enabling the identification of actionable vulnerabilities in residual disease and informing rational design of spatially guided combination therapies.

Contamination of tumour samples with normal epithelium or stromal cells has been shown to systematically bias bulk transcriptomic analyses, leading to misclassification of molecular subtypes (including PAM50) and risk predictors [9, 46–48], misleading differential expression signals[47], and inflated immune or EMT signatures that reflect low purity rather than intrinsic tumour biology[48, 49]. The SMART dataset is uniquely positioned to study transcriptomic profiles of both morphologically and phenotypically distinct regions, owing to the pathologist-guided manual selection of ROIs segmented into cell-type specific AOIs. We utilise these features to select exclusively invasive epithelium, followed by pseudobulking methods to classify each sample according to the PAM50[14], Lehmann [6] and Baylor [15] classifiers and test the prognostic significance of several signatures associated with responses to NACT in TNBC. Notably, this analysis demonstrated that the SMART dataset recapitulates known prognostic signals in pre-NACT TNBCs. Moreover, paired analysis revealed changes in the molecular TNBC subtype following NACT, often reflecting a transition to subtypes associated with poorer prognoses such as Basal-like and BLIS. Together these results highlight the altered state of tumours following NACT and the role of spatial transcriptomics in ensuring tumour purity for subtyping analysis.

Unsupervised clustering of epithelial signatures relating to conserved cell-states revealed seven Epithelial Archetypes (EAs). In line with previous reports[9, 13], several archetypes were defined by the up (EA1) or down (EA2) regulation of cell cycle and CIN, enrichment for immune-responses (EA4) and EMT and ECM-remodelling (EA6). EA1 was shown to be enriched in pre-NACT samples of responders compared to non-responders, concurrent with the observed paradox in TNBC, wherein high levels of proliferation and CIN are associated with both more aggressive tumours and enhanced responses to NACT [50, 51]. Using the SMART dataset to integrate histological and morphological features associated with each EA revealed distinct patterns. Notably, EA2 and EA3 were both characterised by the downregulation of cell cycle and CIN signatures but EA3 was exclusively comprised of histologically ‘normal’ epithelium, unlike EA2 which contained both invasive and noninvasive morphologies. This highlights the importance of spatial diversities in tumour profiling and demonstrates the unique opportunity afforded by the SMART dataset to study transcriptomic profiles from the same tumour along an increasingly ‘malignant’ axis. Additionally, transcriptional signalling in EA6 reflecting polarised states of EMT significantly correlated with opposing patterns of tumour dissemination, highlighting the inherent link between molecular signalling and tumour architecture. Critically, the SMART dataset integrates image-based network analysis with transcriptomic profiles from the same region and slide, enabling these two dimensions to be studied in perfectly matched samples.

Tumour-Immune Microenvironments (TIMEs) were then identified by unsupervised clustering of deconvoluted cell abundances and characterised by distinct compositions of immune subtypes. Notably, we found a cluster of plasma cells that were spatially segregated from other B cell aggregates, in line with previous studies [10, 52], raising the question of origin for infiltrating plasma cells. Additionally, we found that pre-NACT frequencies of TIME1 (enriched for memory CD4+ T cells) were significantly associated non-responders, meanwhile that of TIME2 (enriched for plasma cells) and TIME3 (enriched for B cells, CD4+ T cells and pDCs) were associated with responders. Importantly, we demonstrated that their prognostic significance was independent of sTILs infiltration, highlighting the functional role of B cells co-localised with T cells in promoting NACT responses in TNBC as previously reported [53].

Further analysis of co-localisation patterns identified a subset of TIME3 (B cells, CD4+ T cells and pDCs) AOIs representing immune cell aggregates that exhibited increased expression for a TLS signature when localised around histologically ‘normal’ epithelium from EA3 (downregulated for cell cycle and CIN signatures). Importantly, this co-occurrence was detected exclusively in pre-NACT samples from responders, however, its emergence in residual tumours of non-responders was found to correlate with prolonged overall survival. These results are consistent with previous reports of ‘immature’ TLS’, defined as T and B cell aggregates that lack a central germinal centre [54], around pre-neoplastic hepatic lesions in hepatocellular carcinoma (HC) [42] and raise the question of how ‘normal’ these epithelium in EA3 truly are. We therefore hypothesise that the co-localisation of TIME3 and EA3 is indicative of ‘immature TLS’ around pre-malignant epithelium that is prognostic for NACT responses in the neoadjuvant setting, and confers a clinical benefit to patients with residual disease. Functional studies to elucidate the underlying mechanisms may provide further insight into novel targets to enhance responses to NACT and prolong overall survival. Additionally, we found that when the same co-occurrence of EA and TIMEs appeared in both response groups, for example EA4 co-localised with TIME4, the direction of correlation between tumour-immune cell proximities and the functional activation of immune cells differentiated responders from non-responders in the neoadjuvant setting. These results provide further credence to the notion that spatial variables provide critically important information with clinical relevance and should be considered alongside molecular profiling where possible.

In recent years, pembrolizumab, an immune-checkpoint blockade (ICB) targeting PDL1, in combination with chemotherapy has been approved for patients with early or locally advanced TNBC [55], in both the neoadjuvant and adjuvant setting. We and others have demonstrated the the prognostic significance of increased infiltrating B cells [56] and the presence of TLS [57] as well as the profound influence of chemotherapy on the TME of breast cancer[24]. In line with the immunogenicity of chemotherapy [58], our results indicate the potential of NACT to induce TLS-like immune aggregates that promote enhanced overall survival, concurrent with previous studies that report chemotherapy-induced TLS [59]. Moreover, short-term chemotherapy is shown to have immunostimulatory effects leading to enhanced clinical benefit in response to ICBs [60].

Our study faced several limitations and challenges. Recent advances in ST technologies have facilitated single-cell and subcellular resolution spatially resolved transcriptomics, for example with the Xenium (10x Genomics)[61] and CosMx Spatial Molecular Imager (Bruker/NanoString) [62]. Whilst the GeoMx DSP profiles phenotypically distinct regions defined by morphology markers unique to the cell-type of interest, these AOIs are analysed as mini-bulk compartments comprising of between 20 and several thousand cells. Additionally, staining variations across the tissue, autofluorescence and overlapping cell types can lead to contamination of AOIs, despite optimisation of segmentation techniques. This limitation introduces technical variation between AOIs with vastly different numbers of nuclei and lead to high keratin expression in some immune AOIs. Additionally, the high cost of the GeoMx DSP platform restricted profiling to just several ROIs in the large post-NACT resections. Whilst every effort was made to sample representative regions of the tissue, this approach may not fully capture the spatial heterogeneity of the TME, and important spatio-molecular features present in unsampled areas could have been missed. Moreover, profiling large cohorts on the GeoMx DSP is manually intensive which, combined with the high cost, limits its applications in the clinical setting. Nevertheless, computational approaches to predict spatio-molecular signatures identified from ROIs co-registration with digitised images from routine H&E slides in the SMART dataset has the potential to overcome such challenges and positions our ST dataset as a diagnostic aid.

In conclusion, our study provides an essential resource, the SMART dataset, that will allow researchers to study the determinants of NACT responses and its the longitudinal effects on the TNBC TME at the clinical, histological, morphological, spatio-molecular and protein level. Given the current treatment landscape of TNBC, this study therefore meets a critical gap in the literature that is is essential to our understanding and optimisation of TNBC responses to ICBs in combination with chemotherapy.

## 4 Material and Methods

### 4.1 H&E Staining & Imaging

Tissue sections were first mounted and dried over night at ∼ 37*^◦^*C. Slides were then de-waxed and H&E stained using the Tissue-Tek DRS2000 automatic slide stainer according to the following protocol: xylene for 2 minutes x2, 100% ethanol for 2 minutes x2, 70% ethanol for 2 minutes, tap water for 2 minutes, hematoxylin for 4 minutes, tap water for 1 minute, hydrochloric acid/ethanol for 30 seconds, tap water for 2 minutes, zip water for 30 seconds, eosin for 2 minutes 30 seconds, wash in zip water for 30 seconds, 70% ethanol for 30 seconds, 100% ethanol for 1 minute x2, xylene for 2 minutes x2. Slides were then cover slipped and stored at ∼ 4*^◦^*C.

Slides were imaged for digital pathological analysis on the Hamamatsu NanoZoomer S60 at 20x or 40x magnification, the increased resolution was implemented upon request of the pathologist.

### 4.2 Imaging-Mass Cytometry (IMC)

FFPE or FF tissues were sectioned into 4-micron thick sections and mounted onto standard glass slides. FFPE samples were deparaffinised in xylene and subjected to re-hydration using 1005 ethanol, followed by 95% ethanol and water prior to staining. FF samples were thawed and fixed immediately using 4% paraformaldehyde in PBS.

Following fixation, tissues were blocked and incubated overnight at 4*^◦^*C with a cocktail of metal-conjugated antibodies. Each antibody in the panel was tagged with a unique metal isotope and selected based on target specificity, clone, and catalogue information (Supplementary Table 2). After antibody incubation, samples were treated with the Iridium Cell-ID Intercalator-Ir for nuclear staining and subsequently air-dried.

Slides were then inserted into the Hyperion Imaging System and photographed to aid region selection (Panorama). Regions of 1 mm2 were selected and laser-ablated at 200-Hz frequency at 1*µ*m/pixel resolution. The resulting images were subsequently uploaded to BioImageArchive and are available at https://www.ebi.ac.uk/biostudies/bioimages/studies/S-BIAD2027?key=5aa4522c-16d5-4644-9cb1-e51e92c8f936.

### 4.3 GeoMx DSP Sample Preparation

Formalin-Fixed Paraffin-Embedded (FFPE) tissue sections were dried overnight at 37 °C, baked at 60 °C for one hour, and then processed according to the *RNA Slide Preparation Protocol* (GeoMx DSP Manual Slide Preparation User Manual; MAN-10150-02). Slides underwent deparaffinisation and rehydration (MAN-10150-02), antigen retrieval in 1x Tris-EDTA for 20 min at ∼99 °C, RNA target exposure in 0.1 µg µL^−1^ Proteinase-K for 20 min at 37 °C, post-fix preservation (MAN-10150-02), in situ hybridisation with the human Whole Transcriptome Atlas (WTA) RNA probes overnight at 37 °C (MAN-10150-02), and stringent washes in equal parts 4x Saline Sodium Citrate (SSC) buffer and *>* 99.9% formamide (MAN-10150-02).

Fresh-Frozen (FF) tissues were sectioned and submerged in 10 % Neutral-Buffered Formalin (NBF) overnight to thaw, baked at 60 °C for one hour, and then processed according to Appendix II in the *RNA Slide Preparation Protocol* (GeoMx DSP Manual Slide Preparation User Manual; MAN-10150-02). This was followed by antigen retrieval in 1x Tris-EDTA for 15 min at ∼85 °C, RNA target exposure in 0.1 µg µL^−1^ Proteinase-K for 20 min at 37 °C, in situ hybridisation with the human WTA RNA probes overnight at 37 °C (MAN-10150-02), and stringent washes in equal parts 4x SSC and *>* 99.9% formamide (MAN-10150-02).

Following the stringent washes, all tissues were blocked with Buffer W from the Slide Preparation Kit for 30 min at room temperature and incubated for 1 h at room temperature with each of the following antibody solutions. First, primary anti-CD45 antibody (2B11 with PD7/26) [*Novus Biologicals; #NBP2-34528AF594*] at a concentration of 5 µg µL^−1^ and primary anti-Ki67 antibody (D2H10) [*Cell Signalling; #44092*] at 6 µg µL^−1^. Second, donkey anti-mouse Alexa Fluor^®^ 594 [*Jackson Immuno; #715-586-150*] and donkey anti-rabbit Alexa Fluor^®^ 647 [*Abcam; #ab150075*], both at a 1:600 dilution. Finally, anti-PanCK (AE-1/AE-3) antibody conjugated with Alexa Fluor^®^ 532 [*Novus Biologicals; #NBP2-33200AF532*] at 2 µg µL^−1^, and SYTO 13 Green Fluorescent Nucleic Acid Stain [*Thermo Fisher Scientific; #S7575*] at a 1:1000 dilution. Each incubation was followed by two 5 min washes in 2x SSC buffer to remove unbound compounds.

Slides were stored at 4*^◦^*C in 2x SSC, and regions of interest (ROIs) were selected and segmented into areas of illumination (AOIs) containing PanCK+, CD45+, CD45-/PanCK-, or PanCK-cells, within 21 days, in accordance with NanoString guidelines.

### 4.4 Histological Annotations & Pathologist-Guided ROI Selection

For pre-NACT samples, we selected uniform 250 x 250 *µ*m ROIs across all epithelial cells, where possible, in order the to capture the whole tissue architecture (Fig 2.7, top panel). In some cases, custom ROIs were required to ensure each ROI contained distinct epithelial morphologies. In the post-NACT setting, we selected pathologist-guided ROIs of 660*µ*m diameter at the tumour edge, the tumour centre and any additional regions of histological interest. Each ROI was then segmented into tumour (PanCK+) and immune (CD45+) compartments. In some cases, a “stromal” segment (PanCK-CD45-), tertiary lymphoid structures (TLS) or whole ROIs were also captured. From each cell-pellet, eight 250 x 250 *µ*m ROIs were selected and segmented into PanCK+ segments to act as a control both between in-house batches, and across multiple sites with which we were collaborating.

Each ROI was assigned an epithelial type and tumour region according to the annotations made by a pathologist from the paired H&E images. In brief, these annotations were utilised to ensure selection of morphologically distinct ROIs of normal epithelium (distant or tumour-adjacent), tumour (cancerisation, invasive tumour, ductal carcinoma in situ (DCIS), lymphovascular invasion (LVI), squamous epithelium and cancerisation) and the TME (necrosis, fibrosis, stroma and inflammation). In cases where mixed cytology was unavoidable due to intricate hetero-dispersion of different cellular morphologies, these ROIs were annotated as having ‘mixed’ epithelial. In rare cases where tissue morphology is compromised due to fixation, tearing or ‘crushing’ of cells, these regions were annotated as “Indeterminate”. Additionally, quantification of stromal TILs was conducted on the invasive tumour regions of each tissue, as discrete variables indicating 1%, 5%, 10%, 15%, 20%, 30%, 40%, 50%, 60% or 70% infiltration, as per the guidelines laid out by Salgado *et al.* [63]. Moreover, annotations of tumour region were provided, where applicable, including the tumour edge, centre/core and isolated foci.

### 4.5 RNA Readout

Samples were prepared for sequencing and downstream analysis as per the GeoMx DSP Readout User Manual. Briefly, GeoMx DSP aspirates were dried at 60*^◦^*C for 1 hours to remove any liquid, ligated to sequencing primers and spatial barcodes via PCR, purified with AMPure beads and sequenced on Illumina Next-Seq 500/550 or NovaSeq 2000 platforms at a depth of 100 reads per *µm*^2^.

Raw FASTQ files were processed via the GeoMx NGS Pipeline designed to trim, stitch and deduplicate raw sequencing reads, producing digital count conversion (DCC) files ready for downstream analysis.

### 4.6 Quality Control & Batch Correction

Filtering of the dataset was first performed to remove AOIs that did not meet the following thresholds: minimum 5,000 reads per ROI, ≥ 80% trimmed/stitch/alignment rates, ≥ 50% saturation, ≤ 1, 000, 000 counts in no-template controls (NTCs), ≥ 20 estimated nuclei, and ≥ 1000*µm*^2^ tissue area. Additional QC steps included checking for outliers in nuclei-to-area ratio and area-to-read counts using custom plotting and filtering functions (See Data Availability for related code).

Probe-level QC was then implemented using setBioProbeQCFlags() with filters for low probe signal ratios (*<* 0.1) and Grubbs outlier test (global and local). Outliers were excluded, and modules retained only if ≥ 98% of probes passed QC. Non-specific or anomalously behaving negative control probes were visualised and removed using a custom function, filterNegativeProbes(), retaining only high-performance negative control targets for the Limit of Quantification (LOQ) estimation.

The LOQ was computed using the geometric mean and standard deviation of negative control probes, applying a cut-off exponent of 2 and minimum threshold of 2 counts. Each gene was assessed for detection above LOQ in each AOIs. AOIs with *<* 1% of genes detected were removed, and genes retained only if detected in ≥ 1% of AOIs or designated as negative controls. The final filtered dataset contained high-quality ROIs and reliably detected genes for further analysis. High quality AOIs were then subjected to batch correction using the RUV4 method [64].

Patient- and tissue-level annotations (e.g., BRCA1/2 status, response to NACT, tumour grade, Tumour, Node, Metastasis (TNM) staging, sTIL scores) were assigned according to unique Tissue IDs. Consistency of metadata assignments was validated through grouping and merging checks. Redundant samples (e.g., duplicates or replicates) were removed, and variables were appropriately factorised for downstream modelling and visualisation.

### 4.7 Pseudobulk Analysis

#### 4.7.1 Generation Of Pseudobulk Dataset

To generate pseudobulk dataset, the raw data counts from high quality AOIs identified during previous QC checks were first filtered to contain only those samples of interest (Fig. S4A). Raw counts were then aggregated, shifted by 1, and the remaining probes were used to calculate the NegativeGeoMean. Genes with an average detection below that of the NegativeGeoMean, representative of background noise, were removed and all remaining gene counts were normalised using the counts per million (CPM) method (Fig. S4 B-D). Principal component analysis was used to identify batch effects (Fig. S4D), and outlier samples were removed accordingly.

#### 4.7.2 PAM50 Intrinsic Subtypes

PAM50 classification of pseudobulk gene expression data from PanCK+ AOIs from ROIs surrounding ‘invasive’ tumour cells was performed using the Absolute Intrinsic Molecular Subtyping (AIMS) method [65]. The AIMS algorithm is a rule-based classifier that applies a one-vs-all top-scoring pairs (TSP) approach to gene expression data. Normalised count data were provided as a numeric matrix, and classification was conducted using the applyAIMS() function from the AIMS package, which internally loads a pre-trained model (AIMSmodel) containing discriminatory gene pairs for breast cancer subtyping. To ensure compatibility with the classifier, the input expression matrix was filtered to retain only genes required by the model and to exclude those with excessive zero or negative values, as AIMS operates on expression values greater than zero. The .apply.nbc() function evaluates pairwise gene expression comparisons and assigns each sample to a subtype based on the most frequently satisfied decision rules.

#### 4.7.3 Lehmann TNBC Subtypes

Lehmann classification of pseudobulk gene expression data from PanCK+ AOIs from ROIs surrounding ‘invasive’ tumour cells was performed using the publicly available online tool ‘TNBCType’[66], available at https://cbc.app.vumc.org/tnbc/. A normalised expression matrix was submitted to the tool which initially identified potential ER-positive samples which were subsequently removed from the matrix before resubmitting the remaining samples for analysis. Normalised expression profiles were compared against six predefined subtype centroids using Spearman correlation. A permutation-based approach was implemented to account for variable number of genes defining each subtype and ensure comparability across subtypes. Each sample was classified according to the subtype with the highest correlation coefficient. Samples with low overall correlation (coefficient *<* 0.1 or p-value *>* 0.05), or with similarly high correlations to multiple subtypes (difference between the top two correlations *<* 0.05), were designated as “unstable” (UNS) to reflect ambiguity in subtype assignment.

#### 4.7.4 Baylor Classifier

Pseudobulk gene expression data from PanCK+ AOIs from ROIs surrounding ‘invasive’ tumour cells were compared against reference centroids from the Baylor classification system [15], defined by 80 subtype-specific genes; *DHRS2*, *GABRP*, *AGR2*, *PIP*, *FOXA1*, *PROM1*, *TFF1*, *NAT1*, *BCL11A*, *ESR1*, *FOXC1*, *CA12*, *TFF3*, *SCUBE2*, *SFRP1*, *ERBB4*, *SIDT1*, *PSAT1*, *CHI3L1*, *AR*, *CD36*, *OGN*, *ABCA8*, *CFD*, *IGF1*, *HBB*, *CDH1*, *MEOX2*, *GPX3*, *SCARA5*, *PDK4*, *ENPP2*, *AGTR1*, *LEP*, *LPL*, *DPT*, *TIMP4*, *FHL1*, *SRPX*, *EDNRB*, *SERPINB5*, *SOX10*, *IRX1*, *MIA*, *DSC2*, *TTYH1*, *COL9A3*, *FGL2*, *RARRES3*, *PDE9A*, *BST2*, *PTGER4*, *KCNK5*, *PSMB9*, *HLA-DMA*, *EPHB3*, *IGSF6*, *ST3GAL6*, *RHOH*, *SGPP1*, *CXCL9*, *CXCL11*, *GBP5*, *GZMB*, *LAMP3*, *GBP1*, *ADAMDEC1*, *CCL5*, *SPON1*, *PBK*, *STAT1*, *EZH2*, *PLAT*, *TAP2*, *SLAMF7*, *HERC5*, *SPOCK1*, *TAP1*, *CD2*, *AIM2*. Normalised gene expression counts were subjected to Spearman correlations between each sample and the four Baylor centroids (LAR, MES, BLIS & BLIA). Each sample was assigned to the subtype with the highest correlation coefficient. Correlation matrices and associated p-values were generated using pairwise correlation tests.

#### 4.7.5 Computation of Prognostic TNBC Signatures

Global pseudobulk normalised gene expression counts were used to compute the following signatures according to their original publication. For analysis of spatial variation of each signature score, the same methods were applied to calculate scores from the normalised and batch corrected spatially-resolved expression set previously generated.

The GeparSixto gene signature was calculated as the median of normalised expression values of the following genes: *CXCL9*, *CD8A*, *CCL5*, *CXCL13*, *CR2*, *FOXP3*, *CD80*.

The EMT.6 gene signature was calculated using Gene Set Variation Analysis (GSVA) from the following genes; *LUM*, *SFRP4*, *COL6A3*, *MMP2*, *CXCL12*, *HTRA1*, to calculate the variation in gene expression within each sample rather than across genes. Genes of interest were first ranked according to normalised expression values and a running sum is then computed by iterating over the list of ranked genes, resulting in a final normalised enrichment score (NES).

The ImmuneEight signature was calculated as the mean of normalised expression values of the following genes: *KLRK1*, *JCHAIN*, *CD69*, *CD40LG*, *MS4A1*, *CD1C*, *KLRB1*, *CA4*.

The 20-gene classifier score was calculated as the sum of normalised expression values of the following genes: *GBP1*, *ZNF711*, *IGHV4-59*, *IGHV4-61*, *IGHV4-59*, *IGHV4-61*, *SLC19A2*, *MCM5*, *IGHM*, *LHX2*, *IDO1*, *IGHV4-31*, *IGHM*, *IGHG4*, *IGHG3*, *IGHG1*, *IGHD*, *IGHA2*, *IGHA1*, *IGH@*, *SCNN1A*, *IL2RA*, *CTSS*, *VASH2*, *NUS1P1*, *NUS1*, *MAP9*, *GBP1*, *SLAMF8*, *IL7*, *EFS*, *FBXO*.

The DLDA30 gene signatures was calculated as the mean of normalised expression values of the following genes: *ZNF552*, *THSD4*, *THNSL2*, *SCUBE2*, *RRM2*, *RAMP1*, *PAIP2*, *NKAIN1*, *MGC5370*, *METRN*, *MELK*, *MED13L*, *MAPT*, *MAPT*, *MAPT*, *MAPT*, *KIAA1467*, *JMJD2B*, *IGFBP4*, *GFRA1*, *GAMT*, *ERBB4*, *E2F3*, *CTNND2*, *CA12*, *BTG3*, *BTG3*, *BECN1*, *BBS4*, *AMFR*.

The NACTRINE gene signatures were calculated as the mean of normalised expression values of each gene for the two sets. The following genes were upregulated in patients that achieved pCR, and used to compute the ‘NACTRINE UP’ signature: *FOXC2*, *KCNB1*, *ZBTB16*, *TFF1*, *GRIN2A*, *C5orf38*, *VEGFA*, *HAS1*, *INHBA*, *PGR*, *PLA2G4F*, *BNIP3*, *FGF1*, *DLL4*, *SNAI1*, *ADCY9*, *RAD52*, *HEMK1*. The ‘NACTRINE DOWN’ signature, representing genes that were downregulated in patients that achieved pCR was computed from the following genes: *HDAC2*, *MCM2*, *ALDH1A1*, *FGL2*, *CXADR*, *CXCL9*.

The PAM50 proliferation signature was calculated as the mean of normalised expression values of the following genes: *BIRC5*, *CCNB1*, *CDC20*, *NUF2*, *CEP55*, *NDC80*, *MKI67*, *PTTG1*, *RRM2*, *TYMS*, *UBE2C*.

#### 4.7.6 Outcome Modelling Of Pseudobulk Signatures

To evaluate the prognostic relevance of each signature, a composite data frame was constructed incorporating patient identifiers, treatment response, and multiple signature scores derived from preprocessed pre-NACT data. Treatment response was binarised (Responder = 0, Non-Responder = 1), and scores were normalised where needed. For each treatment cohort, data was filtered to include only patients who received either 1) ECT with or without Carboplatin, 2) ECT only or 3) ECT with Carboplatin. Logistic regression models were then independently fitted for each signature using response as the binary outcome variable. Odds ratios (OR), 95% confidence intervals (CI), and p-values were extracted. The resulting statistics were compiled into a summary table and visualised using a forest plot, highlighting associations between gene expression signatures and treatment outcomes.

### 4.8 Statistical Analysis

Clinical and molecular data from a BC cohort were processed and analysed to explore associations with treatment response and survival outcomes. Samples labelled as “Pellet” were excluded, and patient-level data were aggregated by unique patient identifiers to ensure consistent clinical and molecular features. Numerical variables (e.g., age, stromal tumour-infiltrating lymphocytes) were summarised using descriptive statistics including mean, standard deviation, quartiles, skewness, and kurtosis to assess distributional characteristics and guide subsequent analyses. Normality was tested using the Shapiro-Wilk test, and visualisations such as violin plots and box-plots were employed to compare distributions across responder groups.

For categorical variables (e.g., BRCA mutation status, tumour grade, ethnicity, treatment type), counts and proportions were examined and visualised with bar plots stratified by response status. Pairwise relationships among variables were explored using scatter-plot matrices to detect potential confounders or interactions.

Survival analysis was conducted using both Cox proportional hazards regression and Kaplan-Meier estimation. The Cox model incorporated binary-encoded covariates reflecting age, ethnicity, tumour grade, sTILs levels, treatment group, and response status, allowing simultaneous adjustment for multiple clinicopathological factors. Model assumptions were evaluated using Schoenfeld residuals and log-log survival plots to verify proportional hazards. Kaplan-Meier curves and log-rank tests compared survival distributions between groups defined by key clinical covariates.

All analyses were implemented in Python using the lifelines library. Statistical significance was assessed at the *<* 0.05 level.

### 4.9 Random Forest for treatment response

The Random Forest subset of AOIs was defined to include only those AOIs associated with responder and non-responder patients, and originating from tumour regions annotated as one of the following types: DCIS, Squamous, Vascular Invasion, Lymphovascular Invasion, Invasive, or Cancerisation. This filtering resulted in a total of 3, 658 AOIs, and 1708 ROIs. The number of trees in the Random Forest was set to 10% of the training set size, yielding 365 trees. The parameter *k*, representing the number of top-performing trees retained, was set to *k* = 50. From these, we selected AOIs that appeared in at least 80% of the *k* trees, corresponding to *m* = 40.

The model’s learning curve is designed to determine the number of AOIs required before the learning process slows down. Using the same settings, the learning curve can be fitted to a logarithmic function in a highly significant way, a common behaviour in machine learning setting [67]. The inflection point of this curve indicates when accuracy ceases to improve significantly with additional training data.

Lastly, this model is trained on the transcriptomics profiles of each AOI, with labels representing the patient’s treatment response. However, other labels that allow for the subdivision of spatial transcriptomics data into subsets are also suitable.

### 4.10 Registration between H&E and Spatial Transcriptomics

An automated image registration pipeline was developed to align hematoxylin and eosin (H&E) stained whole slide images (WSIs) with their corresponding multiplex immunofluorescence (mIF) images using the DeeperHistReg framework [68]. Prior to registration, preprocessing steps were applied to isolate tissue-containing regions and standardise resolution. Specifically, tissue regions from each H&E WSI were extracted at 10× magnification using QuPath v0.5.1, and the corresponding mIF images were downsampled to the same 10× resolution using QuPath to ensure consistency in image scale.

The registration pipeline was implemented in Python, incorporating OpenSlide and libvips (via PyVIPS) for efficient WSI handling, and PyTorch for GPU-accelerated computation. Input images were loaded from standardised folder structures with image identifiers. The pipeline supported two operating modes: (1) single-image registration based on specified identifiers, and (2) batch-mode processing for multiple H&E-mIF image pairs. To accelerate computation, both source (H&E) and target (mIF) images were internally downsampled by a factor of two before alignment. Registration was configured to use either rigid or non-rigid transformation modes as provided by DeeperHistReg. Initial alignment relied on SuperPoint keypoint detection and RANSAC-based matching, after which the H&E image was spatially transformed onto the mIF image.

Outputs of the pipeline included the warped H&E image, a dense displacement field for nonrigid alignment, and the 3×3 affine transformation matrix for rigid registration. All transformation parameters and relevant metadata were saved to disk to support downstream analysis and ensure reproducibility. An affine transformation is a linear mapping method that preserves points, straight lines, and planes, but not necessarily angles or distances. Unlike rigid transformations, affine transformations can include: scaling, shearing, rotation, and translation. The transformation is expressed as:

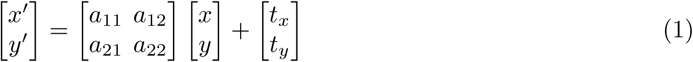

Where, (*x, y*) are the coordinates in the source image, and (*x^′^, y^′^*) are the coordinates in the transformed image. The 2 × 2 matrix

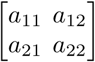

defines the linear transformation, which may include rotation, scaling, and shearing. The vector

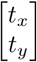

represents the translation component. The affine transformation can also be compactly represented in homogeneous coordinates using a 3 × 3 matrix, as we have done for the downstream analysis, and is written as follows:

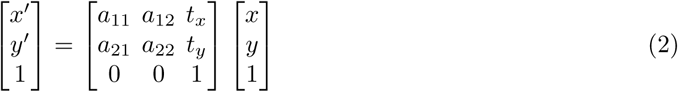

### 4.11 Image Segmentation

Each tissue is associated with a set of ROIs, which are manually adjusted on the GeoMx platform in order to obtain the best possible mask and intensity level for each immunofluorescence channel. The different segments (CD45+, PanCK+, and CD45-/PanCK-) are associated with different pairs of channels (red, green, blue). The CD45+ segment is associated with the blue and red channels, while the PanCK+ segment is associated with the blue and green channels. Finally, artifacts due to the GeoMx process are removed, including the border of the ROI mask.

In addition to cleaning the ROI images, we implemented a quality control (QC) step to retain only those images that behave correctly during the segmentation process performed with StarDist [69]. This QC procedure includes four checks based on the analysis of three channels (red, green, blue), which are derived from the original four fluorescent markers. Although there is no strict correspondence between the channels and the markers, the blue channel is primarily associated with the DNA marker (DAPI).

- area img: The ratio between the area defined by non-zero elements in the blue channel and the total accessible area (i.e., the area defined by the ROI).
- area img std: The standard deviation of the area measurements across the red, green, and blue channels.
- median blue img: The median intensity of the blue channel.
- std blue img: The standard deviation of the intensity values in the blue channel.

Ideally, area img should be high, indicating a large area of cell nuclei within the ROI. A high area img std suggests strong contrast among the different channels, which reflects the presence of distinct cellular compartments (e.g., nuclei, membranes, other markers). Similarly, a high median blue img indicates strong signal from the nuclear marker, and a high std blue img reflects high contrast between nuclei and the surrounding regions of the image.

StarDist is a learning-based approach for segmenting both cells and nuclei in fluorescence microscopy images. In situations involving crowded cells, the method overcomes segmentation errors caused by the poor approximation of bounding boxes by localising cell nuclei through star-convex polygons. Specifically, a convolutional neural network is trained to predict a polygon for the cell instance at each pixel position. These polygons offer a more accurate shape representation compared to bounding boxes and therefore do not require shape refinement.

Once each ROI is segmented, every cell is localised at the centroid of the mask associated with the segmented cell. This process leads to the creation of three different point clouds for each ROI, each associated with a different AOI (segment).

### 4.12 Mean distance between CD45+ and PanCK+ segments

The segmentation of AOIs from the same ROI allows for the computation of the mean distance between cells associated with CD45+ segments (immune cells) and PanCK+ segments (epithelial cells). The segmentation of both cell types within a single ROI yields two point clouds: one corresponding to immune cells and the other to epithelial cells. Since both point clouds are defined within the same coordinate space, we can compute the distance from each epithelial cell to its nearest immune cell. To do this efficiently, we use a k-d tree strategy, which accelerates the nearest-neighbour search and makes the process scalable for large datasets. For each epithelial cell, we calculate the distance to its closest immune cell, and repeat this for all epithelial cells within the ROI. This results in a vector containing the distance from each epithelial cell to its nearest immune cell. Such a vector is computed for each ROI. The mean distance between CD45+ and PanCK+ segments is then defined as the mean value of this vector.

This distance can be computed for each pair of segments across all ROIs, stratified by epithelial archetypes (EA), and further divided into responder and non-responder groups. This enables the assessment of whether the spatial proximity between epithelial and immune cells varies with treatment response and epithelial archetype.

### 4.13 Network Analysis

Network theory provides powerful tools for understanding complex systems by capturing relationships and interactions in a structured and intuitive way. Using the point cloud representations obtained from the cell segmentations, we derive a Euclidean distance matrix, which assesses the distance between every pair of nodes for each AOI. This distance matrix is then filtered using the disparity filter method [70]. This method extracts the relevant backbone of complex multi-scale networks by preserving edges that represent statistically significant deviations from a null model for local edge assignment. In other words, the initial complete weighted network (clique) is filtered into a sparser network containing only significant edges. Unlike alternative methods such as k-nearest neighbours (k-NN) graphs or Delaunay triangulation, which have been used in previous studies and lead to a stereotypical geometry directly determined by their construction method, the disparity filter preserves both local and global geometric properties.

Finally, we study the geometry of these networks and correlate different geometric measures with transcriptomics data. Three different network metrics were used to characterise the geometry of the network associated with each AOI. The average clustering coefficient measures the extent to which nodes connected to a common node are also connected to each other, capturing the network’s clustering tendency. The average shortest path length reflects how close nodes are to one another within the network, providing insight into its overall connectivity and compactness. The degree assortativity quantifies the tendency of nodes to connect with other nodes that have a similar degree, revealing patterns of similarity in node connectivity.

## Data availability

The SMART multimodal dataset is available through PharosAI (https://www.pharosai.co.uk), a Trusted Research Environment (TRE), also referred to as Secure Data Environment. The TRE provided a highly secure computing infrastructure, enabling approved researchers to access and process sensitive datasets in compliance with data protection and confidentiality standards. The PharosAI TRE infrastructure was deployed on-premises using OpenStack, offering a secure, private cloud environment managed by the PharosAI and partners, including the King’s College London e-Research team. All project data within the TRE were encrypted to maintain confidentiality and integrity. Access was strictly controlled through authenticated user permissions tailored to project roles. Additionally, data ingress and egress were tightly regulated through approved protocols to prevent unauthorised transfer or access, ensuring data remained secure throughout the research process. The data can be freely available for research purpose, interesting user have to contact PharosAI using the following email address, in order to have a agreement approved by the Ethics Entity.

## Code availability

The set of tools development with the SMART multimodal dataset are available on the following Github repository: https://github.com/izziWall/SMART

## Acknowledgements

Anthony Baptista, and Anita Grigoriadis acknowledge support from the CRUK City of London Centre Award CTRQQR-2021/100004. Holly Rafique, Pre-Doctoral Fellow NIHR303406, is funded by the NIHR for this research project. Lucy Ryan is funded by the Pathological Society of Great Britain and Ireland and the Jean Shanks Foundation (JSPS CPF 1023 05).

The views expressed in this publication are those of the author(s) and not necessarily those of the NIHR, NHS or the UK Department of Health and Social Care

## Ethics declarations

For archival FFPE samples from the KHP Biobank, ethical approval was obtained from the local research ethics committees (KHP Cancer Biobank REC reference 18/EE/0025). For FF samples obtained through the FORCE clinical trial, ethics were approved by an independent review board (reference nos. 16/LO/1303, NCT03238144).

## References

[1] Bray, F., Laversanne, M., Sung, H., Ferlay, J., Siegel, R.L., Soerjomataram, I., Jemal, A.: Global cancer statistics 2022: GLOBOCAN estimates of incidence and mortality worldwide for 36 cancers in 185 countries. CA: A Cancer Journal for Clinicians 74(3), 229–263 (2024) 10.3322/caac.21834. eprint: https://onlinelibrary.wiley.com/doi/pdf/10.3322/caac.21834. Accessed 2025-03-05

[2] Korde, L.A., Somerfield, M.R., Carey, L.A., Crews, J.R., Denduluri, N., Hwang, E.S., Khan, S.A., Loibl, S., Morris, E.A., Perez, A., Regan, M.M., Spears, P.A., Sudheendra, P.K., Symmans, W.F., Yung, R.L., Harvey, B.E., Hershman, D.L.: Neoadjuvant Chemotherapy, Endocrine Therapy, and Targeted Therapy for Breast Cancer: ASCO Guideline. Journal of Clinical Oncology 39(13), 1485–1505 (2021) 10.1200/JCO.20.03399. Publisher: Wolters Kluwer. Accessed 2025-07-07

[3] Schmid, P., Cortes, J., Dent, R., McArthur, H., Pusztai, L., Kümmel, S., Denkert, C., Park, Y.H., Hui, R., Harbeck, N., Takahashi, M., Im, S.-A., Untch, M., Fasching, P.A., Mouret-Reynier, M.-A., Foukakis, T., Ferreira, M., Cardoso, F., Zhou, X., Karantza, V., Tryfonidis, K., Aktan, G., O’Shaughnessy, J.: Overall Survival with Pembrolizumab in Early-Stage Triple-Negative Breast Cance. New England Journal of Medicine 391(21), 1981–1991 (2024) 10.1056/NEJMoa2409932. Publisher: Massachusetts Medical Society eprint: https://www.nejm.org/doi/pdf/10.1056/NEJMoa2409932. Accessed 2025-07-03

[4] Cortes, J., Rugo, H.S., Cescon, D.W., Im, S.-A., Yusof, M.M., Gallardo, C., Lipatov, O., Barrios, C.H., Perez-Garcia, J., Iwata, H., Masuda, N., Otero, M.T., Gokmen, E., Loi, S., Guo, Z., Zhou, X., Karantza, V., Pan, W., Schmid, P.: Pembrolizumab plus Chemotherapy in Advanced Triple-Negative Breast Cancer. New England Journal of Medicine 387(3), 217–226 (2022) 10.1056/NEJMoa2202809. Publisher: Massachusetts Medical Society eprint: https://www.nejm.org/doi/pdf/10.1056/NEJMoa2202809. Accessed 2025-07-03

[5] Wu, S.Z., Al-Eryani, G., Roden, D.L., Junankar, S., Harvey, K., Andersson, A., Thennavan, A., Wang, C., Torpy, J.R., Bartonicek, N., Wang, T., Larsson, L., Kaczorowski, D., Weisenfeld, N.I., Uytingco, C.R., Chew, J.G., Bent, Z.W., Chan, C.-L., Gnanasambandapillai, V., Dutertre, C.-A., Gluch, L., Hui, M.N., Beith, J., Parker, A., Robbins, E., Segara, D., Cooper, C., Mak, C., Chan, B., Warrier, S., Ginhoux, F., Millar, E., Powell, J.E., Williams, S.R., Liu, X.S., O’Toole, S., Lim, E., Lundeberg, J., Perou, C.M., Swarbrick, A.: A single-cell and spatially resolved atlas of human breast cancers. Nature Genetics 53(9), 1334–1347 (2021) 10.1038/s41588-021-00911-1

[6] Lehmann, B.D., Pietenpol, J.A.: Identification and use of biomarkers in treatment strategies for triple-negative breast cancer subtypes. J Pathol 232(2), 142–50 (2014) 10.1002/path.4280

[7] Barkley, D., Rao, A., Pour, M., França, G.S., Yanai, I.: Cancer cell states and emergent properties of the dynamic tumor system. Genome Research 31(10), 1719–1727 (2021) 10.1101/gr.275308.121. Accessed 2025-05-30

[8] Barkley, D., Moncada, R., Pour, M., Liberman, D.A., Dryg, I., Werba, G., Wang, W., Baron, M., Rao, A., Xia, B., França, G.S., Weil, A., Delair, D.F., Hajdu, C., Lund, A.W., Osman, I., Yanai, I.: Cancer cell states recur across tumor types and form specific interactions with the tumor microenvironment. Nature Genetics 54(8), 1192–1201 (2022) 10.1038/s41588-022-01141-9. Publisher: Nature Publishing Group. Accessed 2025-05-30

[9] Wang, X., Venet, D., Lifrange, F., Larsimont, D., Rediti, M., Stenbeck, L., Dupont, F., Rouas, G., Garcia, A.J., Craciun, L., Buisseret, L., Ignatiadis, M., Carausu, M., Bhalla, N., Masarapu, Y., Villacampa, E.G., Franzén, L., Saarenpää, S., Kvastad, L., Thrane, K., Lundeberg, J., Rothé, F., Sotiriou, C.: Spatial transcriptomics reveals substantial heterogeneity in triple-negative breast cancer with potential clinical implications. Nature Communications 15(1), 10232 (2024) 10.1038/s41467-024-54145-w. Publisher: Nature Publishing Group. Accessed 202502-05

[10] Andersson, A., Larsson, L., Stenbeck, L., Salmén, F., Ehinger, A., Wu, S.Z., Al-Eryani, G., Roden, D., Swarbrick, A., Borg, Frisén, J., Engblom, C., Lundeberg, J.: Spatial deconvolution of HER2-positive breast cancer delineates tumor-associated cell type interactions. Nature Communications 12(1), 6012 (2021) 10.1038/s41467-021-26271-2. Publisher: Nature Publishing Group. Accessed 2025-02-05

[11] Liu, Y.-M., Ge, J.-Y., Chen, Y.-F., Liu, T., Chen, L., Liu, C.-C., Ma, D., Chen, Y.-Y., Cai, Y.-W., Xu, Y.-Y., Shao, Z.-M., Yu, K.-D.: Combined Single-Cell and Spatial Transcriptomics Reveal the Metabolic Evolvement of Breast Cancer during Early Dissemination. Advanced Science 10(6), 2205395 (2023) 10.1002/advs.202205395

[12] Bassiouni, R., Idowu, M.O., Gibbs, L.D., Robila, V., Grizzard, P.J., Webb, M.G., Song, J., Noriega, A., Craig, D.W., Carpten, J.D.: Spatial Transcriptomic Analysis of a Diverse Patient Cohort Reveals a Conserved Architecture in Triple-Negative Breast Cancer. Cancer Research 83(1), 34–48 (2023) 10.1158/0008-5472.can-22-2682

[13] Shiao, S.L., Gouin, K.H., Ing, N., Ho, A., Basho, R., Shah, A., Mebane, R.H., Zitser, D., Martinez, A., Mevises, N.-Y., Ben-Cheikh, B., Henson, R., Mita, M., McAndrew, P., Karlan, S., Giuliano, A., Chung, A., Amersi, F., Dang, C., Richardson, H., Shon, W., Dadmanesh, F., Burnison, M., Mirhadi, A., Zumsteg, Z.S., Choi, R., Davis, M., Lee, J., Rollins, D., Martin, C., Khameneh, N.H., McArthur, H., Knott, S.R.V.: Single-cell and spatial profiling identify three response trajectories to pembrolizumab and radiation therapy in triple negative breast cancer. Cancer Cell 42(1), 70–848 (2024) 10.1016/j.ccell.2023.12.012. Accessed 2025-02-05

[14] Parker, J.S., Mullins, M., Cheang, M.C.U., Leung, S., Voduc, D., Vickery, T., Davies, S., Fauron, C., He, X., Hu, Z., Quackenbush, J.F., Stijleman, I.J., Palazzo, J., Marron, J.S., Nobel, A.B., Mardis, E., Nielsen, T.O., Ellis, M.J., Perou, C.M., Bernard, P.S.: Supervised Risk Predictor of Breast Cancer Based on Intrinsic Subtypes. Journal of Clinical Oncology 27(8), 1160–1167 (2009) 10.1200/JCO.2008.18.1370. Publisher: Wolters Kluwer. Accessed 2025-02-10

[15] Burstein, M.D., Tsimelzon, A., Poage, G.M., Covington, K.R., Contreras, A., Fuqua, S.A.W., Savage, M.I., Osborne, C.K., Hilsenbeck, S.G., Chang, J.C., Mills, G.B., Lau, C.C., Brown, P.H.: Comprehensive Genomic Analysis Identifies Novel Subtypes and Targets of Triple-Negative Breast Cancer. Clinical Cancer Research 21(7), 1688–1698 (2015) 10.1158/1078-0432.Ccr-14-0432

[16] Masuda, H., Baggerly, K.A., Wang, Y., Zhang, Y., Gonzalez-Angulo, A.M., Meric-Bernstam, F., Valero, V., Lehmann, B.D., Pietenpol, J.A., Hortobagyi, G.N., Symmans, W.F., Ueno, N.T.: Differential response to neoadjuvant chemotherapy among 7 triple-negative breast cancer molecular subtypes. Clinical cancer research : an official journal of the American Association for Cancer Research 19(19), 10–115810780432130799 (2013) 10.1158/1078-0432.CCR-13-0799. Accessed 2025-07-05

[17] Gao, G., Wang, Z., Qu, X., Zhang, Z.: Prognostic value of tumor-infiltrating lymphocytes in patients with triple-negative breast cancer: a systematic review and meta-analysis. BMC Cancer 20(1), 179 (2020) 10.1186/s12885-020-6668-z

[18] Loi, S., Salgado, R., Adams, S., Pruneri, G., Francis, P.A., Lacroix-Triki, M., Joensuu, H., Dieci, M.V., Badve, S., Demaria, S., Gray, R., Munzone, E., Drubay, D., Lemonnier, J., Sotiriou, C., Kellokumpu-Lehtinen, P.L., Vingiani, A., Gray, K., André, F., Denkert, C., Piccart, M., Roblin, E., Michiels, S.: Tumor infiltrating lymphocyte stratification of prognostic staging of early-stage triple negative breast cancer. npj Breast Cancer 8(1), 3 (2022) 10.1038/s41523-021-00362-1

[19] Griguolo, G., Paré, L., Miglietta, F., Bottosso, M., Generali, D.G., Frassoldati, A., Musolino, A., Spazzapan, S., Vernaci, G., Giarratano, T., Cappellesso, R., Bisagni, G., Piacentini, F., Tagliafico, E., Cagossi, K., Pinato, C., Schiavi, F., Prat, A., Dieci, M.V., Guarneri, V.: Association of tumor-infiltrating lymphocytes (TILs) and immune signatures with PAM50 intrinsic subtyping hormone receptor-positive (HR+)/HER2-negative (HER2-) early breast cancer (BC): A translational analysis of two multicentric neoadjuvant trials. Journal of Clinical Oncology 42(16 suppl), 558–558 (2024) 10.1200/JCO.2024.42.16 suppl.558. Publisher: Wolters Kluwer. Accessed 2025-04-22

[20] Lehmann, B.D., Jovanović, B., Chen, X., Estrada, M.V., Johnson, K.N., Shyr, Y., Moses, H.L., Sanders, M.E., Pietenpol, J.A.: Refinement of Triple-Negative Breast Cancer Molecular Subtypes: Implications for Neoadjuvant Chemotherapy Selection. PLOS ONE 11(6), 0157368 (2016) 10.1371/journal.pone.0157368. Publisher: Public Library of Science. Accessed 2025-04-22

[21] Harano, K., Wang, Y., Lim, B., Seitz, R.S., Morris, S.W., Bailey, D.B., Hout, D.R., Skelton, R.L., Ring, B.Z., Masuda, H., Rao, A.U.K., Laere, S.V., Bertucci, F., Woodward, W.A., Reuben, J.M., Krishnamurthy, S., Ueno, N.T.: Rates of immune cell infiltration in patients with triplenegative breast cancer by molecular subtype. PLOS ONE 13(10), 0204513 (2018) 10.1371/journal.pone.0204513. Publisher: Public Library of Science. Accessed 2025-07-04

[22] Kim, C., Gao, R., Sei, E., Brandt, R., Hartman, J., Hatschek, T., Crosetto, N., Foukakis, T., Navin, N.: Chemoresistance Evolution in Triple-Negative Breast Cancer Delineated by Single Cell Sequencing. Cell 173(4), 879–89313 (2018) 10.1016/j.cell.2018.03.041. Accessed 2025-02-05

[23] Masuda, H., Harano, K., Miura, S., Wang, Y., Hirota, Y., Harada, O., Jolly, M.K., Matsunaga, Y., Lim, B., Wood, A.L., Parinyanitikul, N., Jin Lee, H., Gong, G., George, J.T., Levine, H., Lee, J., Wang, X., Lucci, A., Rao, A., Schweitzer, B.L., Lawrence, O.R., Seitz, R.S., Morris, S.W., Hout, D.R., Nakamura, S., Krishnamurthy, S., Ueno, N.T.: Changes in Triple-Negative Breast Cancer Molecular Subtypes in Patients Without Pathologic Complete Response After Neoadjuvant Systemic Chemotherapy. JCO precision oncology 6, 2000368 (2022) 10.1200/PO.20.00368

[24] Wall, I., Boulat, V., Shah, A., Blenman, K.R.M., Wu, Y., Alberts, E., Calado, D.P., Salgado, R., Grigoriadis, A.: Leveraging the Dynamic Immune Environment Triad in Patients with Breast Cancer: Tumour, Lymph Node, and Peripheral Blood. Cancers (Basel) 14(18) (2022) 10.3390/cancers14184505

[25] Martín, M., Prat, A., Rodríguez-Lescure, Caballero, R., Ebbert, M.T.W., Munárriz, B., Ruiz-Borrego, M., Bastien, R.R.L., Crespo, C., Davis, C., Rodríguez, C.A., López-Vega, J.M., Furió, V., García, A.M., Casas, M., Ellis, M.J., Berry, D.A., Pitcher, B.N., Harris, L., Ruiz, A., Winer, E., Hudis, C., Stijleman, I.J., Tuck, D.P., Carrasco, E., Perou, C.M., Bernard, P.S.: PAM50 proliferation score as a predictor of weekly paclitaxel benefit in breast cancer. Breast Cancer Research and Treatment 138(2), 457–466 (2013) 10.1007/s10549-013-2416-2. Accessed 2025-02-10

[26] Gluz, O., Kolberg-Liedtke, C., Prat, A., Christgen, M., Gebauer, D., Kates, R., Paré, L., Grischke, E.-M., Forstbauer, H., Braun, M., Warm, M., Hackmann, J., Uleer, C., Aktas, B., Schumacher, C., Kuemmel, S., Wuerstlein, R., Pelz, E., Nitz, U., Kreipe, H.H., Harbeck, N.: Efficacy of deescalated chemotherapy according to PAM50 subtypes, immune and proliferation genes in triple-negative early breast cancer: Primary translational analysis of the WSG-ADAPT-TN trial. International Journal of Cancer 146(1), 262–271 (2020) 10.1002/ijc.32488. eprint: https://onlinelibrary.wiley.com/doi/pdf/10.1002/ijc.32488. Accessed 2025-02-11

[27] Edlund, K., Madjar, K., Lebrecht, A., Aktas, B., Pilch, H., Hoffmann, G., Hofmann, M., Kolberg, H.-C., Boehm, D., Battista, M., Seehase, M., Stewen, K., Gebhard, S., Cadenas, C., Marchan, R., Brenner, W., Hasenburg, A., Koelbl, H., Solbach, C., Gehrmann, M., Tanner, B., Weber, K.E., Loibl, S., Sachinidis, A., Rahnenführer, J., Schmidt, M., Hengstler, J.G.: Gene Expression–Based Prediction of Neoadjuvant Chemotherapy Response in Early Breast Cancer: Results of the Prospective Multicenter EXPRESSION Trial. Clinical Cancer Research 27(8), 2148–2158 (2021) 10.1158/1078-0432.CCR-20-2662. Accessed 2025-02-10

[28] Tseng, L.-M., Huang, C.-C., Tsai, Y.-F., Chen, J.-L., Chao, T.-C., Lai, J.-I., Lien, P.-J., Lin, Y.-S., Feng, C.-J., Chen, Y.-J., Chiu, J.-H., Hsu, C.-Y., Liu, C.-Y.: Correlation of an immune-related 8-gene panel with pathologic response to neoadjuvant chemotherapy in patients with primary breast cancers. Translational Oncology 38, 101782 (2023) 10.1016/j.tranon.2023.101782. Accessed 2025-02-11

[29] Filho, O.M., Stover, D.G., Asad, S., Ansell, P.J., Watson, M., Loibl, S., Geyer, C.E. Jr, Bae, J., Collier, K., Cherian, M., O’Shaughnessy, J., Untch, M., Rugo, H.S., Huober, J.B., Golshan, M., Sikov, W.M., Minckwitz, G., Rastogi, P., Maag, D., Wolmark, N., Denkert, C., Symmans, W.F.: Association of Immunophenotype With Pathologic Complete Response to Neoadjuvant Chemotherapy for Triple-Negative Breast Cancer: A Secondary Analysis of the BrighTNess Phase 3 Randomized Clinical Trial. JAMA Oncology 7(4), 603–608 (2021) 10.1001/jamaoncol.2020.7310. Accessed 2025-02-10

[30] Freitas, A.J.A., Nunes, C.R., Mano, M.S., Causin, R.L., Santana, I.V.V., Oliveira, M.A., Calfa, S., Silveira, H.C.S., Pádua Souza, C., Marques, M.M.C.: Gene expression alterations predict the pathological complete response in triple-negative breast cancer exploratory analysis of the NACATRINE trial. Scientific Reports 13(1), 21411 (2023) 10.1038/s41598-023-48657-6. Publisher: Nature Publishing Group. Accessed 2025-02-10

[31] Wei, L.Y., Zhang, X.J., Wang, L., Hu, L.N., Zhang, X.D., Li, L., Gao, J.N.: A Six-Epithelial–Mesenchymal Transition Gene Signature May Predict Metastasis of Triple-Negative Breast Cancer. OncoTargets and therapy 13, 6497–6509 (2020) 10.2147/OTT.S256818. Accessed 2025-02-10

[32] Hess, K.R., Anderson, K., Symmans, W.F., Valero, V., Ibrahim, N., Mejia, J.A., Booser, D., Theriault, R.L., Buzdar, A.U., Dempsey, P.J., Rouzier, R., Sneige, N., Ross, J.S., Vidaurre, T., Gómez, H.L., Hortobagyi, G.N., Pusztai, L.: Pharmacogenomic predictor of sensitivity to preoperative chemotherapy with paclitaxel and fluorouracil, doxorubicin, and cyclophosphamide in breast cancer. Journal of Clinical Oncology: Official Journal of the American Society of Clinical Oncology 24(26), 4236–4244 (2006) 10.1200/JCO.2006.05.6861

[33] Pathak, N., Sharma, A., Elavarasi, A., Sankar, J., Deo, S.V.S., Sharma, D.N., Mathur, S., Kumar, S., Prasad, C.P., Kumar, A., Batra, A.: Moment of truth-adding carboplatin to neoadjuvant/adjuvant chemotherapy in triple negative breast cancer improves overall survival: An individual participant data and trial-level Meta-analysis. The Breast : Official Journal of the European Society of Mastology 64, 7–18 (2022) 10.1016/j.breast.2022.04.006. Accessed 2025-04-28

[34] Pirrotta, S., Masatti, L., Corrá, A., Pedrini, F., Esposito, G., Martini, P., Risso, D., Romualdi, C., Calura, E.: signifinder enables the identification of tumor cell states and cancer expression signatures in bulk, single-cell and spatial transcriptomic data. bioRxiv, 2023–0307530940 (2023) 10.1101/2023.03.07.530940. Accessed 2025-02-21

[35] Acerbi, I., Cassereau, L., Dean, I., Shi, Q., Au, A., Park, C., Chen, Y.Y., Liphardt, J., Hwang, E.S., Weaver, V.M.: Human breast cancer invasion and aggression correlates with ECM stiffening and immune cell infiltration. Integrative Biology 7(10), 1120–1134 (2015) 10.1039/c5ib00040h. Accessed 2025-07-15

[36] Danaher, P., Kim, Y., Nelson, B., Griswold, M., Yang, Z., Piazza, E., Beechem, J.M.: Advances in mixed cell deconvolution enable quantification of cell types in spatial transcriptomic data. Nature Communications 13(1), 385 (2022) 10.1038/s41467-022-28020-5. Publisher: Nature Publishing Group. Accessed 2025-02-24

[37] Loi, S., Sirtaine, N., Piette, F., Salgado, R., Viale, G., Van Eenoo, F., Rouas, G., Francis, P., Crown, J.P.A., Hitre, E., Azambuja, E., Quinaux, E., Di Leo, A., Michiels, S., Piccart, M.J., Sotiriou, C.: Prognostic and Predictive Value of Tumor-Infiltrating Lymphocytes in a Phase III Randomized Adjuvant Breast Cancer Trial in Node-Positive Breast Cancer Comparing the Addition of Docetaxel to Doxorubicin With Doxorubicin-Based Chemotherapy: BIG 02-98. Journal of Clinical Oncology 31(7), 860–867 (2013) 10.1200/JCO.2011.41.0902. Publisher: Wolters Kluwer. Accessed 2025-07-25

[38] Garaud, S., Buisseret, L., Solinas, C., Gu-Trantien, C., Wind, A., Eynden, G., Naveaux, C., Lodewyckx, J.N., Boisson, A., Duvillier, H., Craciun, L., Ameye, L., Veys, I., Paesmans, M., Larsimont, D., Piccart-Gebhart, M., Willard-Gallo, K.: Tumor infiltrating B-cells signal functional humoral immune responses in breast cancer. JCI Insight 5 (2019) 10.1172/jci. insight.129641

[39] Boisséere-Michot, F., Chateau, M.-C., Thézenas, S., Lafont, V., Crapez, E., Sharma, P., Bobrie, A., Roger, P., Guiu, S., Jacot, W.: Prognostic value of tertiary lymphoid structures in triple-negative breast cancer: integrated analysis with the tumor microenvironment and clinicopathological features. Frontiers in Immunology 15, 1507371 (2024) 10.3389/fimmu.2024.1507371

[40] Schumacher, T.N., Thommen, D.S.: Tertiary lymphoid structures in cancer. Science 375(6576), 9419 (2022) 10.1126/science.abf9419. Publisher: American Association for the Advancement of Science. Accessed 2025-06-05

[41] Sautés-Fridman, C., Dimberg, A., Verma, V.: Editorial: Tertiary Lymphoid Structures: From Basic Biology to Translational Impact in Cancer. Frontiers in Immunology 13, 870862 (2022) 10.3389/fimmu.2022.870862. Accessed 2025-07-28

[42] Meylan, M., Petitprez, F., Lacroix, L., Di Tommaso, L., Roncalli, M., Bougoüin, A., Laurent, A., Amaddeo, G., Sommacale, D., Regnault, H., Derman, J., Charpy, C., Lafdil, F., Pawlotsky, J.-M., Sautés-Fridman, C., Fridman, W.H., Calderaro, J.: Early Hepatic Lesions Display Immature Tertiary Lymphoid Structures and Show Elevated Expression of Immune Inhibitory and Immunosuppressive Molecules. Clinical Cancer Research 26(16), 4381–4389 (2020) 10.1158/1078-0432.CCR-19-2929. Accessed 2025-07-25

[43] Oshi, M., Asaoka, M., Tokumaru, Y., Yan, L., Matsuyama, R., Ishikawa, T., Endo, I., Takabe, K.: CD8 T Cell Score as a Prognostic Biomarker for Triple Negative Breast Cancer. International Journal of Molecular Sciences 21(18), 6968 (2020) 10.3390/ijms21186968. Accessed 2025-06-19

[44] Chen, H.-H., Yu, Y.-R., Hsiao, Y.-L., Chen, S.-H., Lee, C.-K.: Plasmacytoid Dendritic Cells Enhance T-Independent B Cell Response through a p38 MAPK-STAT1 Axis. Journal of Immunology (Baltimore, Md.: 1950) 211(4), 576–590 (2023) 10.4049/jimmunol.2200210

[45] Sluis, R.M., García-Rodríguez, J.L., Nielsen, I.H., Gris-Oliver, A., Becker, J., Costa, B., Chaudhry, M.Z., Werner, M., Laustsen, A., Pedersen, J.G., Gammelgaard, K.R., Mogensen, T.H., Kalinke, U., Cicin-Sain, L., Bak, R.O., Kristensen, L.S., Jakobsen, M.R.: Distinctive CD8+ T cell activation by antigen-presenting plasmacytoid dendritic cells compared to conventional dendritic cells. Cell Reports 44(3), 115413 (2025) 10.1016/j.celrep.2025.115413. Accessed 2025-06-19

[46] Elloumi, F., Hu, Z., Li, Y., Parker, J.S., Gulley, M.L., Amos, K.D., Troester, M.A.: Systematic Bias in Genomic Classification Due to Contaminating Non-neoplastic Tissue in Breast Tumor Samples. BMC Medical Genomics 4(1), 54 (2011) 10.1186/1755-8794-4-54. Accessed 2025-08-25

[47] Aran, D., Sirota, M., Butte, A.J.: Systematic pan-cancer analysis of tumour purity. Nature Communications 6, 8971 (2015) 10.1038/ncomms9971. Accessed 2025-08-25

[48] Koudijs, K.K.M., Scheltinga, A.G.T., Böhringer, S., Schimmel, K.J.M., Guchelaar, H.-J.: The impact of estimated tumour purity on gene expression-based drug repositioning of Clear Cell Renal Cell Carcinoma samples. Scientific Reports 9(1), 2495 (2019) 10.1038/s41598-019-39891-y. Publisher: Nature Publishing Group. Accessed 2025-08-25

[49] Rhee, J.-K., Jung, Y.C., Kim, K.R., Yoo, J., Kim, J., Lee, Y.-J., Ko, Y.H., Lee, H.H., Cho, B.C., Kim, T.-M.: Impact of Tumor Purity on Immune Gene Expression and Clustering Analyses across Multiple Cancer Types. Cancer Immunology Research 6(1), 87–97 (2018) 10.1158/2326-6066.CIR-17-0201. Accessed 2025-08-25

[50] Birkbak, N.J., Eklund, A.C., Li, Q., McClelland, S.E., Endesfelder, D., Tan, P., Tan, I.B., Richardson, A.L., Szallasi, Z., Swanton, C.: Paradoxical Relationship between Chromosomal Instability and Survival Outcome in Cancer. Cancer Research 71(10), 3447–3452 (2011) 10.1158/0008-5472.CAN-10-3667. Accessed 2025-06-02

[51] Keam, B., Im, S.-A., Lee, K.-H., Han, S.-W., Oh, D.-Y., Kim, J.H., Lee, S.-H., Han, W., Kim, D.-W., Kim, T.-Y., Park, I.A., Noh, D.-Y., Heo, D.S., Bang, Y.-J.: Ki-67 can be used for further classification of triple negative breast cancer into two subtypes with different response and prognosis. Breast Cancer Research : BCR 13(2), 22 (2011) 10.1186/bcr2834. Accessed 2025-06-02

[52] Fitzsimons, E., Qian, D., Enica, A., Thakkar, K., Augustine, M., Gamble, S., Reading, J.L., Litchfield, K.: A pan-cancer single-cell RNA-seq atlas of intratumoral B cells. Cancer Cell 42(10), 1784–17974 (2024) 10.1016/j.ccell.2024.09.011. Accessed 2025-07-21

[53] Kuroda, H., Jamiyan, T., Yamaguchi, R., Kakumoto, A., Abe, A., Harada, O., Masunaga, A.: Tumor-infiltrating B cells and T cells correlate with postoperative prognosis in triplenegative carcinoma of the breast. BMC Cancer 21(1), 286 (2021) 10.1186/s12885-021-08009-x. Accessed 2025-04-28

[54] Chen, Y., Wu, Y., Yan, G., Zhang, G.: Tertiary lymphoid structures in cancer: maturation and induction. Frontiers in Immunology 15, 1369626 (2024) 10.3389/fimmu.2024.1369626. Accessed 2025-07-25

[55] 1 Recommendations | Pembrolizumab for neoadjuvant and adjuvant treatment of triple-negative early or locally advanced breast cancer | Guidance | NICE. Publisher: NICE (2022). https://www.nice.org.uk/guidance/ta851/chapter/1-Recommendations Accessed 2025-08-24

[56] García-Martínez, E., Gil, G.L., Benito, A.C., González-Billalabeitia, E., Conesa, M.A.V., García, T.G., García-Garre, E., Vicente, V., De La Penã, F.A.: Tumor-infiltrating immune cell profiles and their change after neoadjuvant chemotherapy predict response and prognosis of breast cancer. Breast Cancer Research 16(6) (2014) 10.1186/s13058-014-0488-5

[57] Cabrita, R., Lauss, M., Sanna, A., Donia, M., Skaarup Larsen, M., Mitra, S., Johansson, I., Phung, B., Harbst, K., Vallon-Christersson, J., Schoiack, A., Lövgren, K., Warren, S., Jirström, K., Olsson, H., Pietras, K., Ingvar, C., Isaksson, K., Schadendorf, D., Schmidt, H., Bastholt, L., Carneiro, A., Wargo, J.A., Svane, I.M., Jönsson, G.: Tertiary lymphoid structures improve immunotherapy and survival in melanoma. Nature 577(7791), 561–565 (2020) 10.1038/s41586-019-1914-8. Publisher: Nature Publishing Group. Accessed 2025-07-09

[58] Zhang, J., Pan, S., Jian, C., Hao, L., Dong, J., Sun, Q., Jin, H., Han, X.: Immunostimulatory Properties of Chemotherapy in Breast Cancer: From Immunogenic Modulation Mechanisms to Clinical Practice. Frontiers in Immunology 12, 819405 (2022) 10.3389/fimmu.2021.819405. Accessed 2025-08-25

[59] Morcrette, G., Hirsch, T.Z., Badour, E., Pilet, J., Caruso, S., Calderaro, J., Martin, Y., Imbeaud, S., Letouzé, E., Rebouissou, S., Branchereau, S., Taque, S., Chardot, C., Guettier, C., Scoazec, J.-Y., Fabre, M., Brugiéres, L., Zucman-Rossi, J.: APC germline hepatoblastomas demonstrate cisplatin-induced intratumor tertiary lymphoid structures. OncoImmunology 8(6), 1583547 (2019) 10.1080/2162402X.2019.1583547. Publisher: Taylor & Francis eprint: 10.1080/2162402X.2019.1583547. Accessed 2025-08-25

[60] Voorwerk, L., Slagter, M., Horlings, H.M., Sikorska, K., Vijver, K.K., Maaker, M., Nederlof, I., Kluin, R.J.C., Warren, S., Ong, S., Wiersma, T.G., Russell, N.S., Lalezari, F., Schouten, P.C., Bakker, N.A.M., Ketelaars, S.L.C., Peters, D., Lange, C.A.H., Werkhoven, E., Tinteren, H., Mandjes, I.A.M., Kemper, I., Onderwater, S., Chalabi, M., Wilgenhof, S., Haanen, J., Salgado, R., Visser, K.E., Sonke, G.S., Wessels, L.F.A., Linn, S.C., Schumacher, T.N., Blank, C.U., Kok, M.: Immune induction strategies in metastatic triple-negative breast cancer to enhance the sensitivity to PD-1 blockade: the TONIC trial. Nat Med 25(6), 920–928 (2019) 10.1038/s41591-019-0432-4

[61] Marco Salas, S., Kuemmerle, L.B., Mattsson-Langseth, C., Tismeyer, S., Avenel, C., Hu, T., Rehman, H., Grillo, M., Czarnewski, P., Helgadottir, S., Tiklova, K., Andersson, A., Rafati, N., Chatzinikolaou, M., Theis, F.J., Luecken, M.D., Wählby, C., Ishaque, N., Nilsson, M.: Optimizing Xenium In Situ data utility by quality assessment and best-practice analysis workflows. Nature Methods 22(4), 813–823 (2025) 10.1038/s41592-025-02617-2. Publisher: Nature Publishing Group. Accessed 2025-07-01

[62] He, S., Bhatt, R., Brown, C., Brown, E.A., Buhr, D.L., Chantranuvatana, K., Danaher, P., Dunaway, D., Garrison, R.G., Geiss, G., Gregory, M.T., Hoang, M.L., Khafizov, R., Killingbeck, E.E., Kim, D., Kim, T.K., Kim, Y., Klock, A., Korukonda, M., Kutchma, A., Lewis, Z.R., Liang, Y., Nelson, J.S., Ong, G.T., Perillo, E.P., Phan, J.C., Phan-Everson, T., Piazza, E., Rane, T., Reitz, Z., Rhodes, M., Rosenbloom, A., Ross, D., Sato, H., Wardhani, A.W., Williams-Wietzikoski, C.A., Wu, L., Beechem, J.M.: High-plex imaging of RNA and proteins at subcellular resolution in fixed tissue by spatial molecular imaging. Nature Biotechnology 40(12), 1794–1806 (2022) 10.1038/s41587-022-01483-z. Publisher: Nature Publishing Group. Accessed 2025-04-28

[63] Salgado, R., Denkert, C., Demaria, S., Sirtaine, N., Klauschen, F., Pruneri, G., Wienert, S., Eynden, G., Baehner, F.L., Penault-Llorca, F., Perez, E.A., Thompson, E.A., Symmans, W.F., Richardson, A.L., Brock, J., Criscitiello, C., Bailey, H., Ignatiadis, M., Floris, G., Sparano, J., Kos, Z., Nielsen, T., Rimm, D.L., Allison, K.H., Reis-Filho, J.S., Loibl, S., Sotiriou, C., Viale, G., Badve, S., Adams, S., Willard-Gallo, K., Loi, S.: The evaluation of tumor-infiltrating lymphocytes (TILs) in breast cancer: recommendations by an International TILs Working Group 2014. Annals of Oncology 26(2), 259–271 (2015) 10.1093/annonc/mdu450. Accessed 2025-03-20

[64] Gagnon-Bartsch, J.A., Jacob, L., Speed, T.P.: Removing Unwanted Variation from High Dimensional Data with Negative Controls

[65] Paquet, E.R., Hallett, M.T.: Absolute assignment of breast cancer intrinsic molecular subtype. Journal of the National Cancer Institute 107(1), 357 (2015) 10.1093/jnci/dju357

[66] Chen, X., Li, J., Gray, W.H., Lehmann, B.D., Bauer, J.A., Shyr, Y., Pietenpol, J.A.: TNBCtype: A Subtyping Tool for Triple-Negative Breast Cancer. Cancer Informatics 11, 9983 (2012) 10.4137/CIN.S9983. Publisher: SAGE Publications Ltd STM. Accessed 2025-02-10

[67] Rajput, D., Wang, W.-J., Chen, C.-C.: Evaluation of a decided sample size in machine learning applications. BMC Bioinformatics 24(1), 48 (2023) 10.1186/s12859-023-05156-9

[68] Wodzinski, M., Müller, H.: DeepHistReg: Unsupervised Deep Learning Registration Framework for Differently Stained Histology Samples. Computer Methods and Programs in Biomedicine 198, 105799 (2021) 10.1016/j.cmpb.2020.105799. Accessed 2025-04-24

[69] Schmidt, U., Weigert, M., Broaddus, C., Myers, G.: Cell Detection with Star-Convex Polygons. In: Frangi, A.F., Schnabel, J.A., Davatzikos, C., Alberola-López, C., Fichtinger, G. (eds.) Medical Image Computing and Computer Assisted Intervention – MICCAI 2018, pp. 265–273. Springer, Cham (2018)

[70] Serrano, M., Boguñá, M., Vespignani, A.: Extracting the multiscale backbone of complex weighted networks. Proceedings of the National Academy of Sciences 106(16), 6483–6488 (2009) 10.1073/pnas.0808904106. Publisher: Proceedings of the National Academy of Sciences. Accessed 2025-02-11

