## Supplementary Information for "SMART: A Spatio-Molecular Atlas of Response Trajectories in Triple-Negative Breast Cancer"

| **Variable** | **All Patients** (n = 96) | **Responders** (n = 51) | **Non-Responders** (n = 43) | **Other** (n = 2) | **Fisher's p-value** |
|---|---|---|---|---|---|
| **Age** | | | | | |
| < 50 | 58 (60.4%) | 34 (66.7%) | 23 (53.5%) | 1 (50.0%) | 0.12 |
| ≥ 50 | 38 (39.6%) | 17 (33.3%) | 20 (46.5%) | 1 (50.0%) | 0.72 |
| **BRCA1** | | | | | |
| Wild Type | 40 (41.7%) | 21 (41.2%) | 19 (44.2%) | 0(0.0%) | 0.86 |
| Mutation | 7 (7.3%) | 6 (11.8%) | 0 (0.0%) | 1(50.0%) | 0.03 |
| VUS | 1 (1.0%) | 0 (0.0%) | 1 (2.3%) | 0(0.0%) | 1.00 |
| Missing Data | 48 (50.0%) | 24 (47.1%) | 23 (53.5%) | 1(50.0%) | NA |
| **Ethnicity** | | | | | |
| White | 35 (36.5%) | 17 (33.3%) | 17 (39.5%) | 1(50.0%) | 1.00 |
| Black | 24 (25.0%) | 13 (25.5%) | 10 (23.3%) | 1 (50.0%) | 0.66 |
| Asian | 7 (7.3%) | 6 (11.8%) | 1 (2.3%) | 0 (0.0%) | 0.12 |
| Mixed | 5 (5.2%) | 3 (5.9%) | 2 (4.7%) | 0 (0.0%) | 1.00 |
| Other | 4 (4.2%) | 2 (3.9%) | 2 (4.7%) | 0 (0.0%) | NA |
| Not Stated | 20 (20.8%) | 10 (19.6%) | 10 (23.3%) | 0 (0.0%) | 1.00 |
| Missing Data | 1 (1.0%) | 0 (0.0%) | 1 (2.3%) | 0 (0.0%) | NA |
| **NACT Group** | | | | | |
| EC-T | 27 (28.1%) | 12 (23.5%) | 15 (34.9%) | 0 (0.0%) | 0.68 |
| EC-T + Carboplatin | 46 (47.9%) | 33 (64.7%) | 13 (30.2%) | 0 (0.0%) | 0.00 |
| Other | 23 (24.0%) | 6 (11.8%) | 15 (34.9%) | 2 (100.0%) | NA |
| **RCB Group** | | | | | |
| 0 (pCR) | 45 (46.9%) | 45 (88.2%) | 0 (0.0%) | 0 (0.0%) | 0.0 |
| I | 6 (6.2%) | 6 (11.8%) | 0 (0.0%) | 0 (0.0%) | 0.03 |
| II | 20 (20.8%) | 0 (0.0%) | 20 (46.5%) | 0 (0.0%) | 0.0 |
| III | 13 (13.5%) | 0 (0.0%) | 13 (30.2%) | 0 (0.0%) | 0.0 |
| Other | 2 (2.1%) | 0 (0.0%) | 0 (0.0%) | 2 (100.0%) | NA |
| Missing Data | 10 (10.4%) | 0 (0.0%) | 10 (23.3%) | 0 (0.0%) | NA |
| **Grade** | | | | | |
| Grade 2 | 5 (5.2%) | 2 (3.9%) | 3 (7.0%) | 0 (0.0%) | 1.00 |
| Grade 3 | 77 (80.2%) | 44 (86.3%) | 31 (72.1%) | 2 (100.0%) | 0.08 |
| Missing Data | 14 (14.6%) | 5 (9.8%) | 9 (20.9%) | 0 (0.0%) | NA |
| **sTILs** | | | | | |
| Low (<30%) | 39 (40.6%) | 20 (39.2%) | 18 (41.9%) | 1 (50.0%) | 0.86 |
| High (≥30%) | 27 (28.1%) | 16 (31.4%) | 10 (23.3%) | 1 (50.0%) | 0.29 |
| Missing Data | 30 (31.2%) | 15 (29.4%) | 15 (34.9%) | 0 (0.0%) | NA |

**Table S1** Baseline clinico-pathological features of the patient cohort (n = 96), stratified by treatment response status. The final column reports Fisher's exact test p-values for each comparison; "NA" indicates that a sub-category comparison was not applicable or not reported.

| Tag # | Tag | Catalogue Number | Target | Clone |
|---|---|---|---|---|
| 141 | Pr | 3141018D | CD38 | EPR4106 |
| 142 | Nd | 3142013D | EGFR | D38B1 |
| 143 | Nd | 3143026D | p53 | DO-7 |
| 144 | Nd | 3144025D | CD14 | EPR3653 |
| 145 | Nd | 3145015D | T-bet/TBX21 | D6N8B |
| 146 | Nd | 3146020D | CD16 | EPR16784 |
| 147 | Sm | 3147021D | CD163 | EDHu-1 |
| 148 | Nd | 3148020D | Pan-Keratin | C11 |
| 149 | Sm | 3149028D | CD11b | EPR1344 |
| 150 | Nd | 3150031D | PD-L1 | E1L3N |
| 151 | Eu | 3151021D | CD107a/LAMP1 | H4A3 |
| 152 | Sm | 3152018D | CD45 | D9M8I |
| 153 | Eu | 3153029D | CD44 | IM7 |
| 154 | Sm | 3154024D | CD366 | D5D5R |
| 155 | Gd | 3155018D | FoxP3 | PCH101 |
| 156 | Gd | 3156033D | CD4 | EPR6855 |
| 158 | Gd | 3158029D | E-Cadherin | 24E10 |
| 159 | Tb | 3159035D | CD68 | KP1 |
| 160 | Gd | ab7856 | Mouse monoclonal [CR3/43] to HLA DR + DP + DQ | CR3/43 |
| 161 | Dy | 3161029D | CD20 | H1 |
| 162 | Dy | 3162034D | CD8a | C8/144B |
| 163 | Dy | 3163028D | VEGF | G153-694 |
| 164 | Dy | EH12.2H7 | Purified CD279 (PD-1) (Maxpar® Ready) Antibody | EH12.2H7 |
| 165 | Ho | 3165032D | $\beta$-Catenin | D13A1 |
| 166 | Er | 3166030D | B7-H4 | H74 |
| 167 | Er | 3167021D | Granzyme B | EPR20129-217 |
| 168 | Er | 3168022D | Ki-67 | B56 |
| 169 | Tm | 3169023D | Collagen Type I | Polyclonal |
| 170 | Er | 3170019D | CD3 | Polyclonal, C-Terminal |
| 171 | Yb | 3171024D | CD27 | EPR8569 |
| 172 | Yb | 3172031D | PD-L2 | D7U8C |
| 173 | Yb | 3173016D | CD45RO | UCHL1 |
| 174 | Yb | ab7817 | Mouse monoclonal [1A4] to alpha smooth muscle Actin | 1A4 |
| 175 | Lu | ab193555 | Rabbit monoclonal [EPR3776] to Vimentin - Cytoskeleton Marker | EPR3776 |
| 176 | Yb | NB600-562SS | CD-31 | JC/70A |

**Table S2** Imaging-Mass Cytometry (IMC) Antibody Panel used in the study. Each antibody is tagged with a unique metal isotope and includes catalogue information, target antigen, and clone specification.

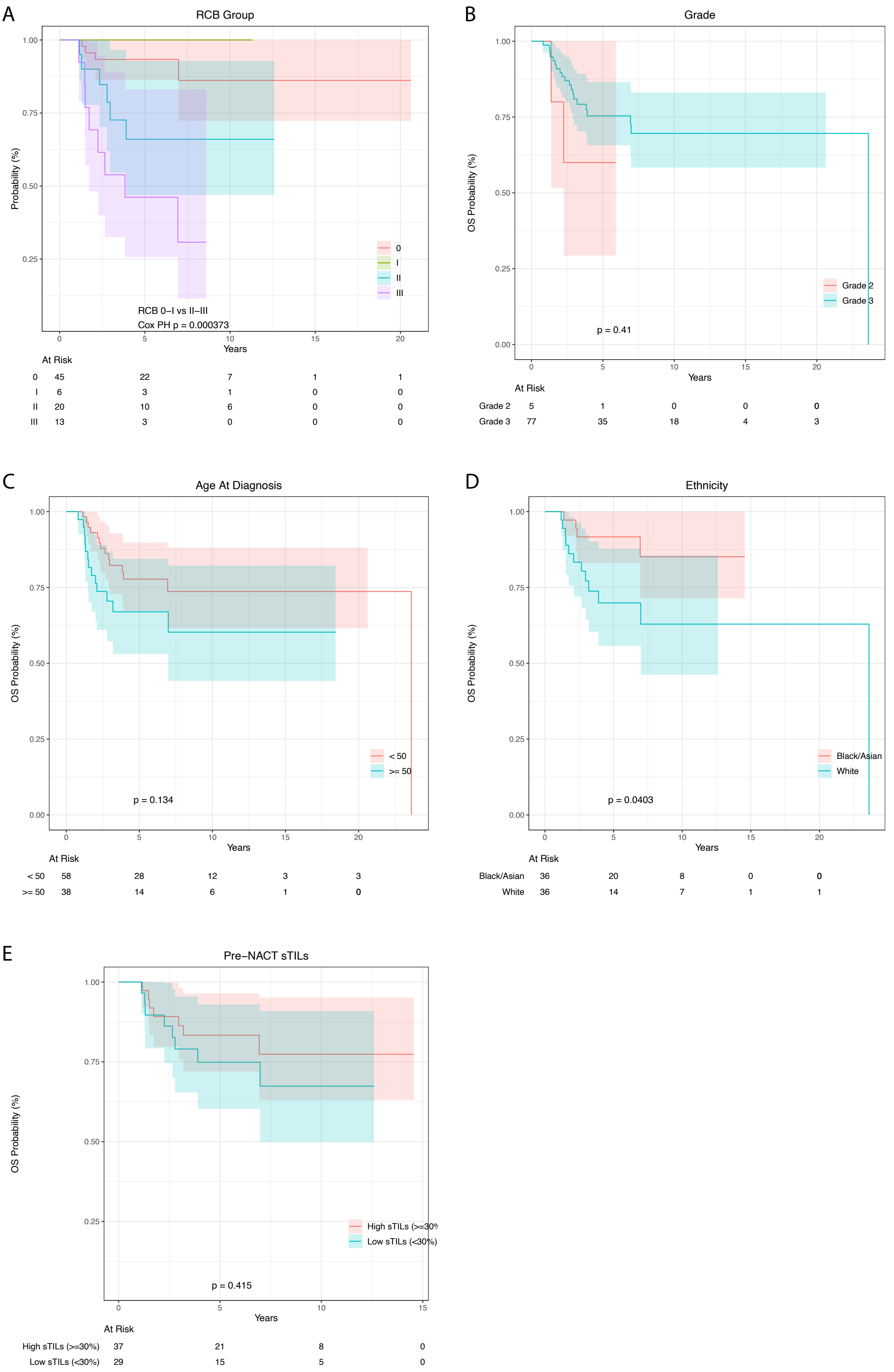


**Fig. S1** Kaplein-Mayer Survival Curves For Clinicopathological Characteristics: Kaplan–Meier log-log survival estimates for overall survival from time of diagnosis in months are shown for **A)** stromal tumour-infiltrating lymphocytes (sTILs) analysed by a pathologist in diagnostic FFPE core needle biopsies comparing high ($\geq$ 30%) versus low ($<$ 30%) sTILs, **B)** histological grade analysed by a pathologist in diagnostic FFPE core needle biopsies comparing high (Grade 3) versus low (Grades 1-2) tumours, **C)** age at diagnosis comparing young ($\leq$ 50 years) versus old ($>$ 50 years) and **D)** self-identified ethnicity comparing white versus non-white patients. Shaded areas represent 95% confidence intervals. Total patient numbers per group are given in the key within the plot. P-values were calculated using the log-rank test with significance considered as p-value $>$ 0.05.

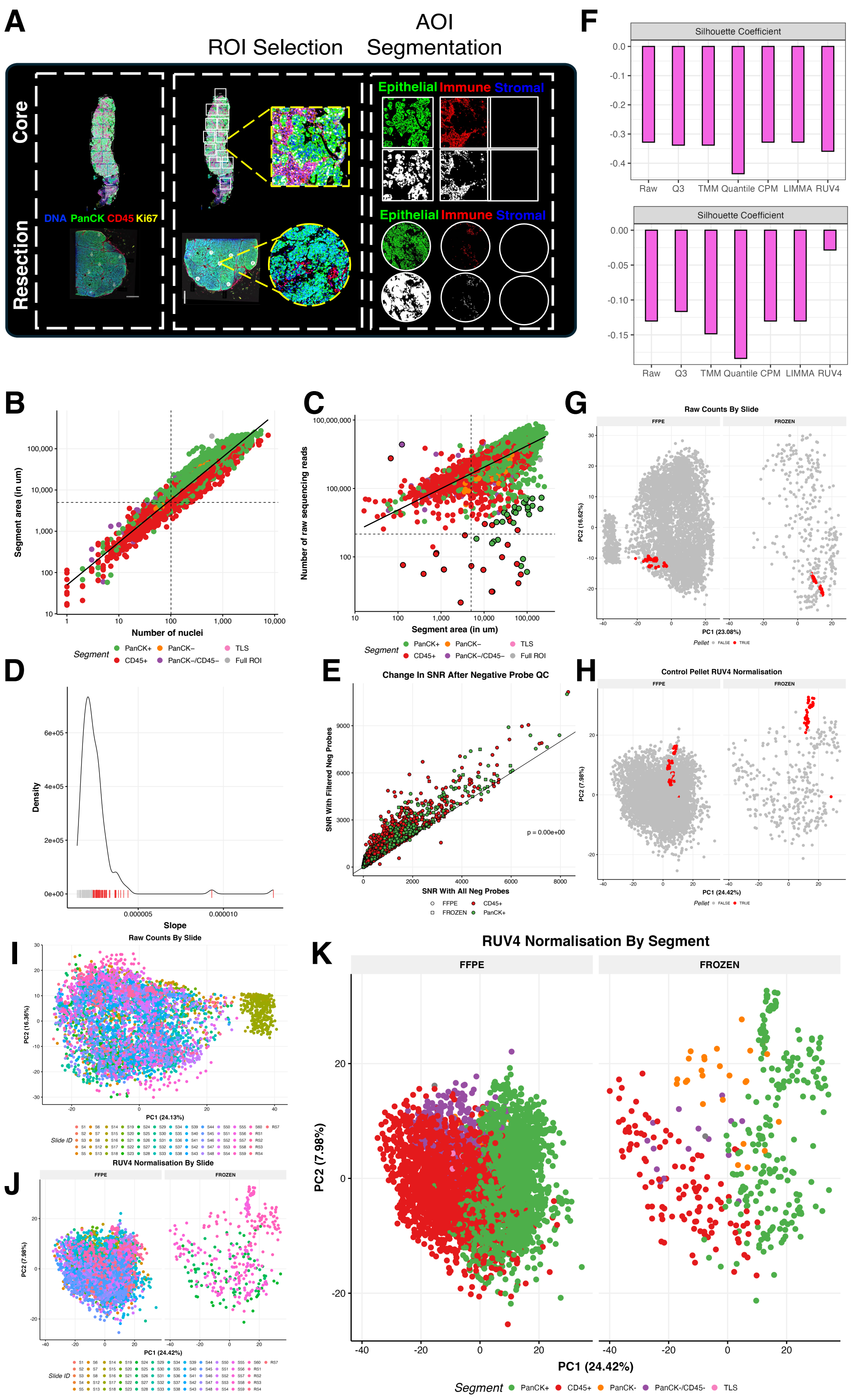


**Fig. S2** Generation & Quality Control Of GeoMx DSP Dataset: **A)**Representative fluorescent images captured on the GeoMx DSP of pre-NACT core biopsies (top) and post-NACT surgical resections (bottom) stained for DNA (blue), PanCK (green), CD45 (red) and Ki67 (yellow). Regions of interest (ROI) were selected with the gridding tool (top) or circle tool (bottom) and further segmented into areas of illumination (AOI) based on expression of fluorescent markers to capture epithelial (DNA+ PanCK+ CD45-, green), immune (DNA+ PanCK- CD45+, red) and stromal (DNA+ PanCK- CD45-, blue) cells. Dotplots indicating the correlation and line of best fit between **B)** the number of nuclei and raw sequencing reads and **C)** the area and raw sequencing reads per AOI. Dots are coloured according to the AOI (segment) and outliers are outlined with a solid black line. **D)** Density plot showing the slope of all negative control probes. Below the slope, outlier probes to be removed are indicated in red. **E)**Dotplot depicting the change in signal-to-noise ratio (SNR) per AOI observed before and after the negative probe QC. Dots above the line at y=0 indicate those with an increased SNR, whilst those below had reduced SNR. Dots are coloured according to AOI. **F)** Silhouette scores illustrating the degree of separation between clusters based on technical (top) and biological (bottom) variation. Higher silhouette scores indicate better-defined, more distinct clusters. Effective methods should result in low silhouette scores for technical variation and high scores for biological variation. Principal-Component Analysis (PCA) plots for **G)** raw and **H)** batch-corrected counts to highlight clustering of AOIs obtained from control pellets in red for both FFPE and Frozen tissues. Principal-Component Analysis (PCA) plots for **I)** raw and batch-corrected counts coloured by batch (Slide ID) **J)** and AOI/segment (**K**) to demonstrate efficacy of RUV4 to remove unwanted batch effects

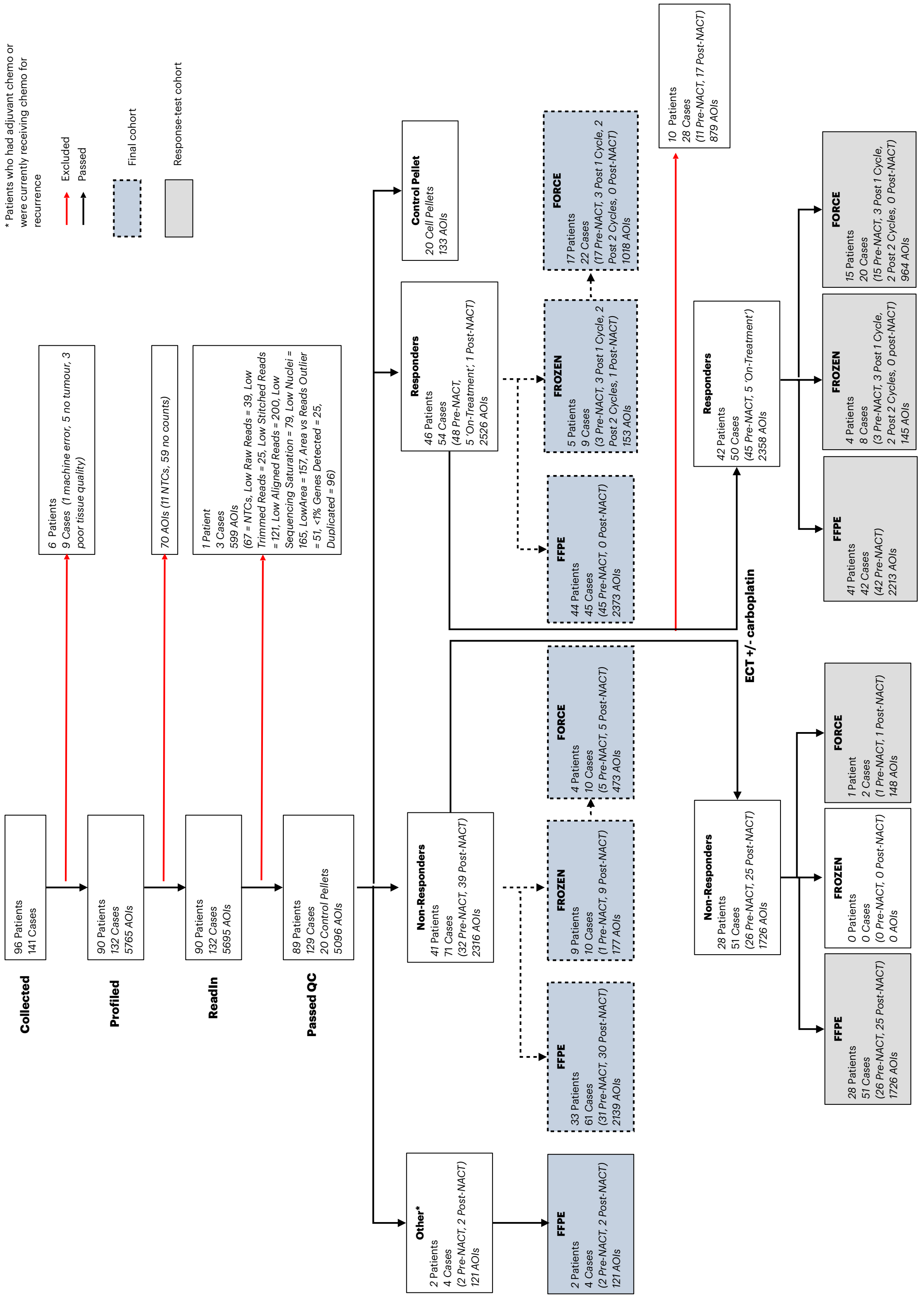


**Fig. S3 Consort Diagram indicating the flow of samples collected, profiled, read into R for downstream analysis, passed QC and stratified by cohort & clinical parameters.** Black solid arrows indicate the direction of flow for samples passing each QC stage. Red solid arrows indicate samples lost at each QC stage. Black dotted arrows indicate stratification of patients by tissue preservation method and for FROZEN blocks, those obtained via the FORCE clinical trial. Blue shaded boxes indicate the samples that passed all stages of QC and grey shaded boxes indicate those that were included in outcome analysis that had comparable treatment regimens.

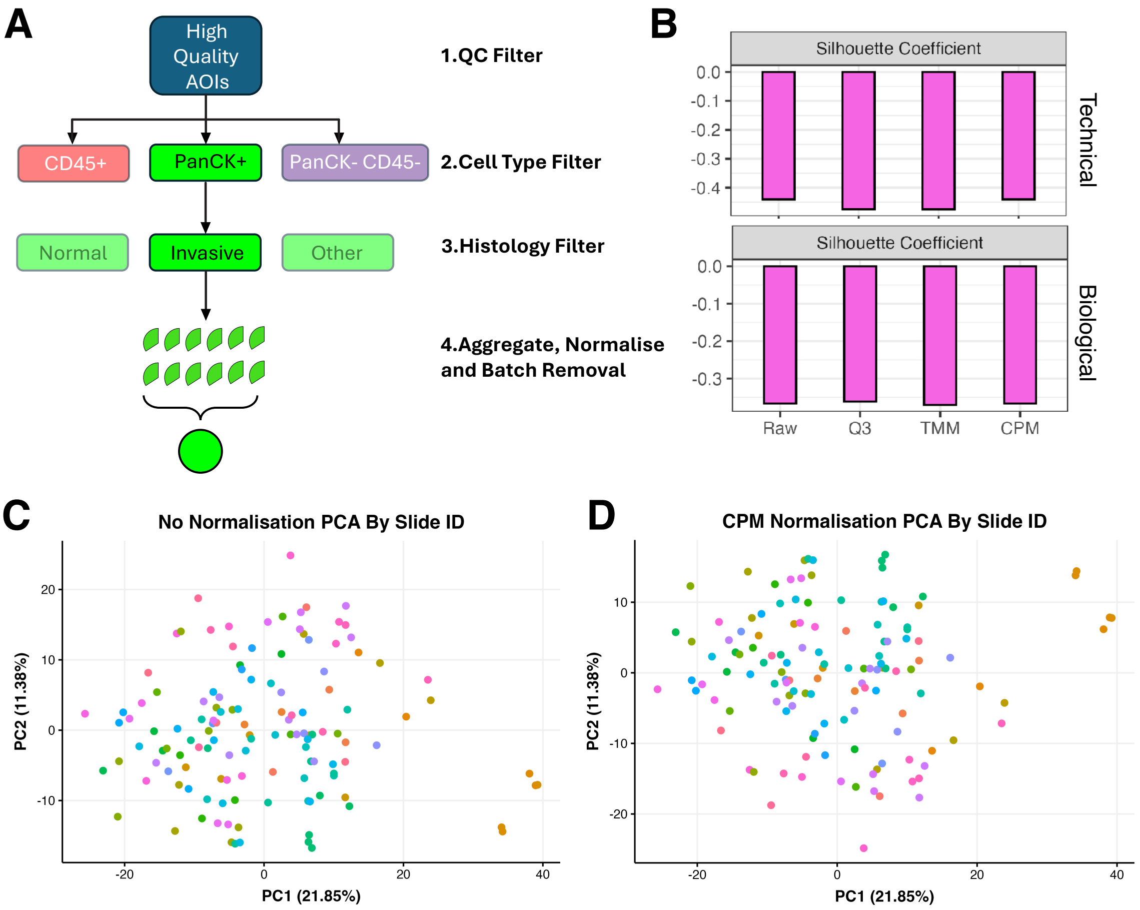


**Fig. S4** Generation & Normalisation Of Pseudobulk Datasets: **A)** Schematic representing the workflow to select samples of interest based on cell-type/AOI and histological annotations for pseudobulking. **B**Silhouette scores illustrating the degree of separation between clusters in pseudobulk data based on technical (top) and biological (bottom) variation. Higher silhouette scores indicate better-defined, more distinct clusters. Effective methods should result in low silhouette scores for technical variation and high scores for biological variation. Principal-Component Analysis (PCA) plots for **C)** raw and **D)** batch-corrected counts coloured by batch (Slide ID) to identify the presence of batch effects before and after correction.

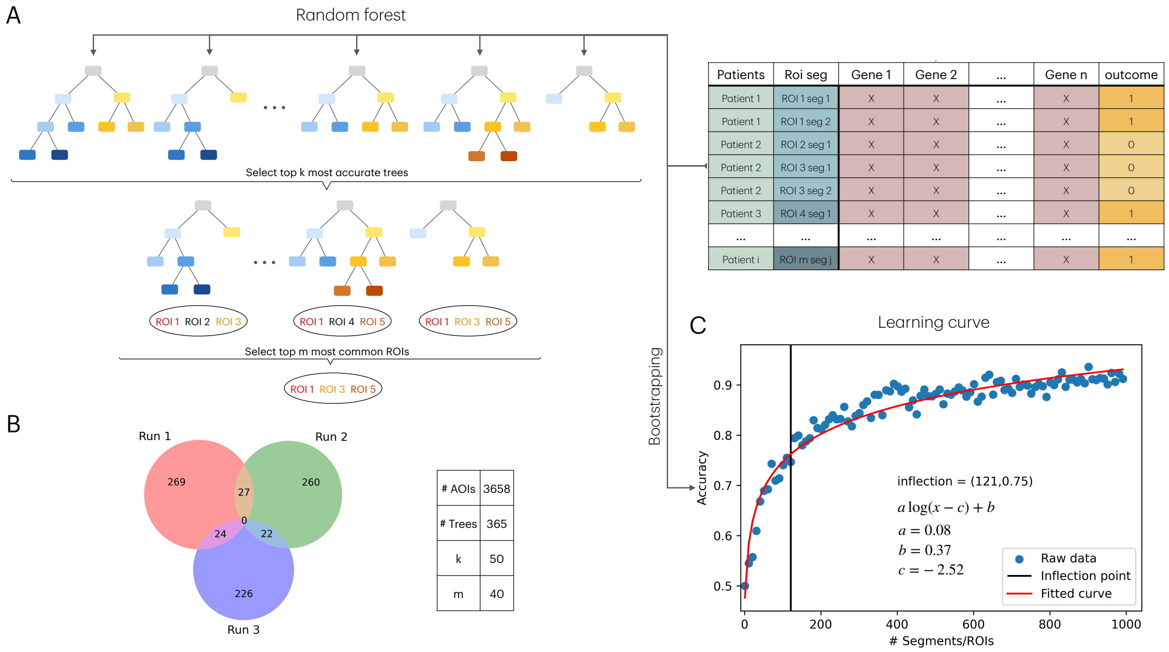


**Fig. S5 Random Forest strategy to define the predictive power of ROIs: A)** A Random Forest model is trained on transcriptomics data (shown in the table on the right) associated with each ROI, using treatment response labels corresponding to each patient. To identify the most predictive ROIs, we first select the top $k$ decision trees with the highest classification accuracy. From this subset of trees, we retain only those ROIs that appear in at least $m$ of the $k$ trees. Notably, the Random Forest is designed to avoid separating segments from the same ROI, as the batch correlation between segments cannot be disentangled. **B)** Venn diagram of three independent runs of the Random Forest strategy described in A, showing that each run yields a different subset of predictive ROIs. the table on the right display the key parameters of the Random Forest strategy. **C)** The Random Forest strategy described in A is used to determine the number of ROIs required to reach an accuracy plateau. Each data point represents performance on a bootstrapped subset of ROIs. Experimental results (blue points) are overlaid with a logarithmic fit (red curve), a common approach to model learning saturation. Parameters of the fitted logarithmic function are displayed within the plot. The vertical black line indicates the inflection point, marking where the increase in accuracy begins to slow.

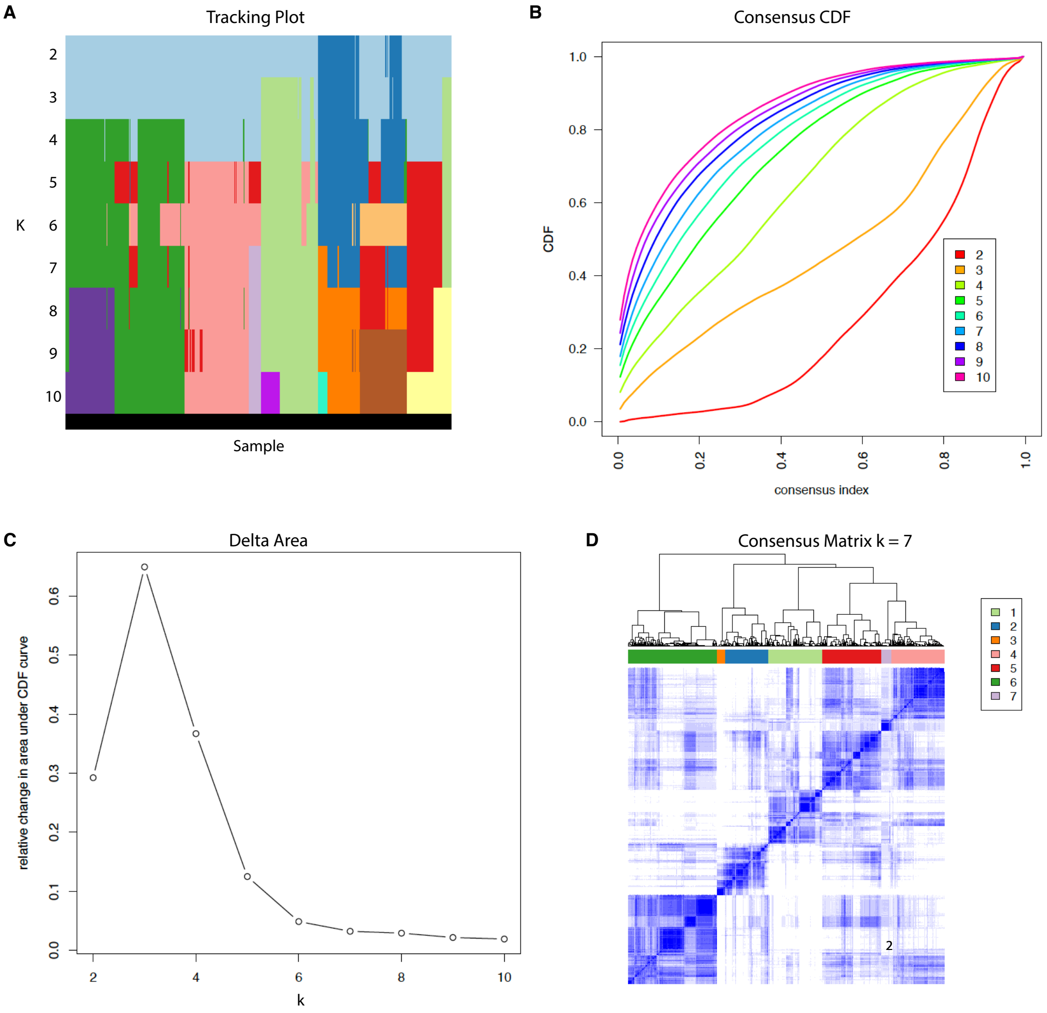


**Fig. S6** Consensus Cluster Plus Identifies The Most Stable Number Of Clusters Based On Signature Expression In PanCK+ AOIs. **A)** Consensus clustering track plot showing sample assignment (x-axis) across increasing cluster (k) values (y-axis). Each row corresponds to a specific number of clusters (k), and each column represents an individual sample. Colour bands indicate cluster membership, allowing visualization of how sample assignments evolve as k increases. Stable clustering is reflected by consistent colour patterns across successive k values. **B)** Consensus cumulative distribution function (CDF) plot showing the cumulative density (y-axis) of the consensus index (x-axis) for each number of clusters (k). Each curve represents a different k value, coloured accordingly. The shape of the CDF curves provides insight into cluster stability, with flatter curves indicating greater consensus among sample assignments. **C)** Elbow plot showing the relative change in area under the cumulative distribution function (CDF) curve (y-axis) for each number of clusters (k, x-axis). The plot illustrates the delta area, which reflects the incremental gain in clustering stability. A noticeable decrease in the delta area indicates diminishing returns with increasing k, aiding in the selection of an optimal number of clusters. **D)** Consensus matrix heatmap for k = 7, showing the pairwise consensus values between samples based on resampling across clustering iterations. Each cell represents the proportion of times two samples were clustered together. Higher consensus values (closer to 1, shown in blue) indicate stronger clustering agreement. The clear block-diagonal structure reflects well-defined and stable clusters.

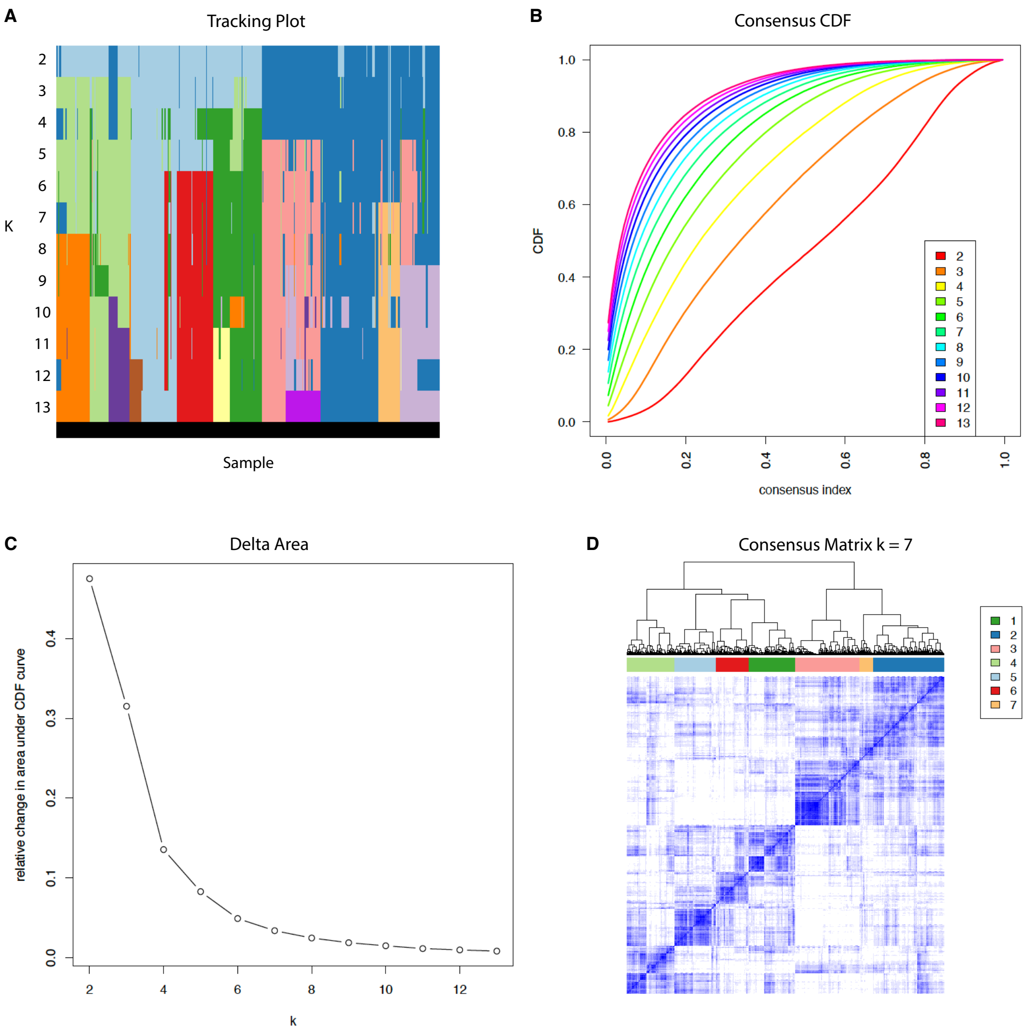


**Fig. S7** Consensus Cluster Plus Identifies The Most Stable Number Of Clusters Based On Relative Cell-Type Proportions in CD45+ AOIs **A)** Consensus clustering track plot showing sample assignment (x-axis) across increasing cluster (k) values (y-axis). Each row corresponds to a specific number of clusters (k), and each column represents an individual sample. Colour bands indicate cluster membership, allowing visualization of how sample assignments evolve as k increases. Stable clustering is reflected by consistent colour patterns across successive k values. **B)** Consensus cumulative distribution function (CDF) plot showing the cumulative density (y-axis) of the consensus index (x-axis) for each number of clusters (k). Each curve represents a different k value, coloured accordingly. The shape of the CDF curves provides insight into cluster stability, with flatter curves indicating greater consensus among sample assignments. **C)** Elbow plot showing the relative change in area under the cumulative distribution function (CDF) curve (y-axis) for each number of clusters (k, x-axis). The plot illustrates the delta area, which reflects the incremental gain in clustering stability. A noticeable decrease in the delta area indicates diminishing returns with increasing k, aiding in the selection of an optimal number of clusters. **D)** Consensus matrix heatmap for k = 7, showing the pairwise consensus values between samples based on resampling across clustering iterations. Each cell represents the proportion of times two samples were clustered together. Higher consensus values (closer to 1, shown in blue) indicate stronger clustering agreement. The clear block-diagonal structure reflects well-defined and stable clusters.

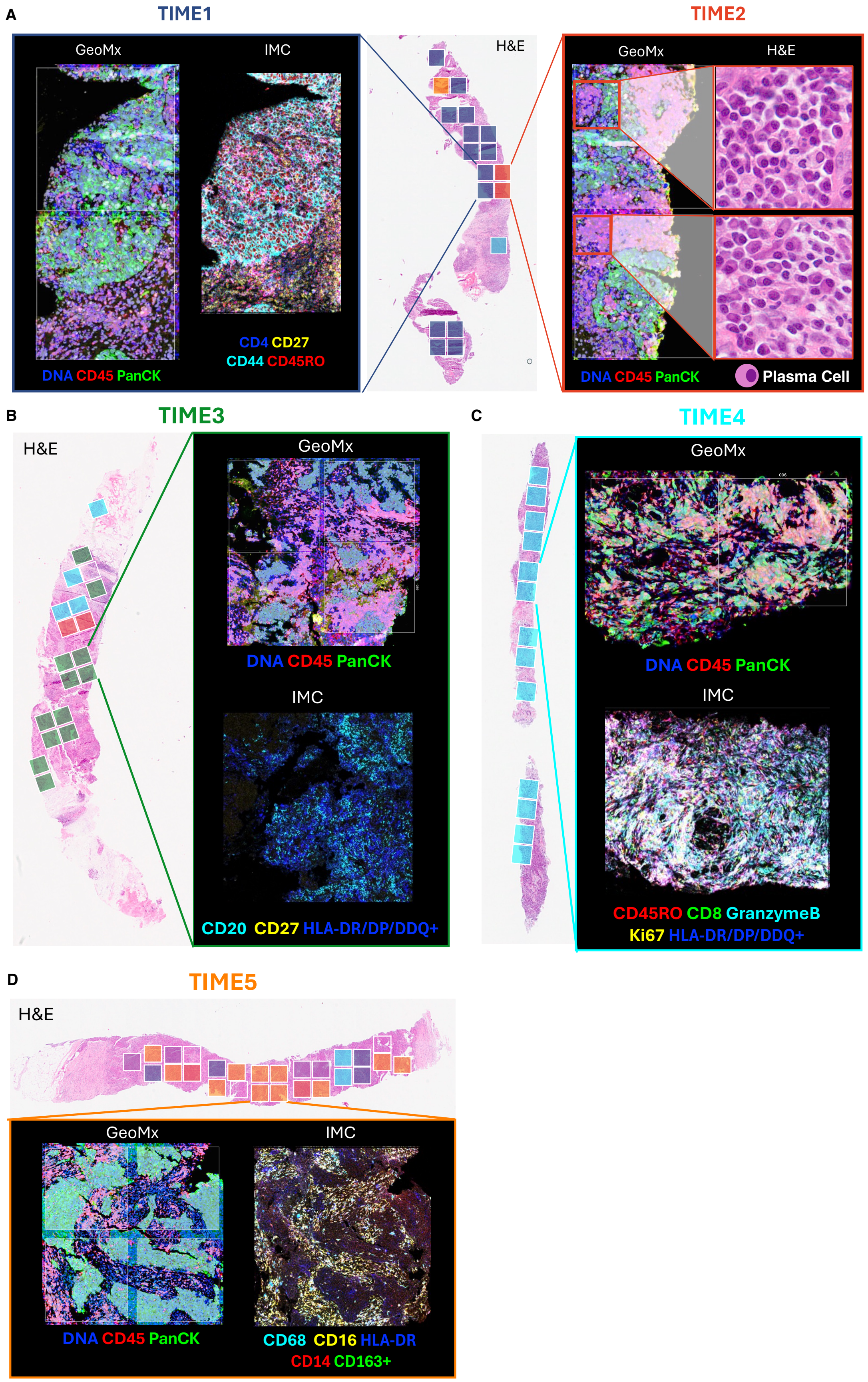


**Fig. S8** Imaging-Mass Cytometry (IMC) Indicates The Presence of Major TIME Cell Types In Corresponding ROIs from Serial Sections **A)** Schematic diagram indicates the selection of IMC ROIs to correspond with those selected on the GeoMx DSP from serial sections. **B-D)** Representative IF images obtained from GeoMx DSP ROIs and the corresponding IMC ROIs displaying protein expression of the relevant targets identifying major phenotypic cell types characteristic of TIME1-5. The marker for plasma cells (CD38 – TIME2) failed to produce a detectable signal. However H&E images were able to validate the presence of plasma cells, identified by their prominent pale perinuclear area in the cytoplasm (black arrow), corresponding to the Golgi apparatus [1], and an eccentric nucleus (red arrow).

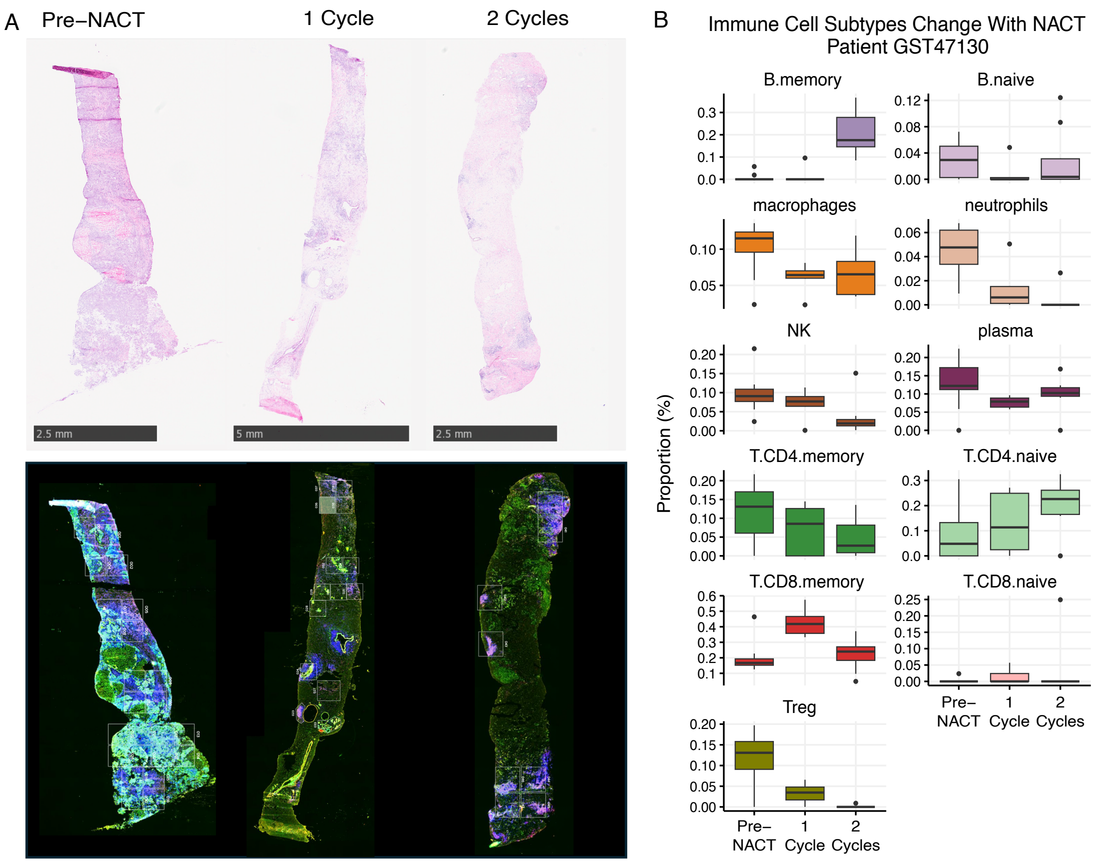


**Fig. S9** Case Study Of Changes In Immune Cell Subtypes After Consecutive Cycles Of NACT In One Patient: **A)** Representative H&E (top) and fluorescent GeoMx DSP (bottom) images of patient-matched samples obtained pre-NACT (left), after one cycle of NACT (middle) and after two cycles of NACT (right). **B)** Boxplots depicting the relative proportions of immune cell subtypes before and after consecutive cycles of NACT.

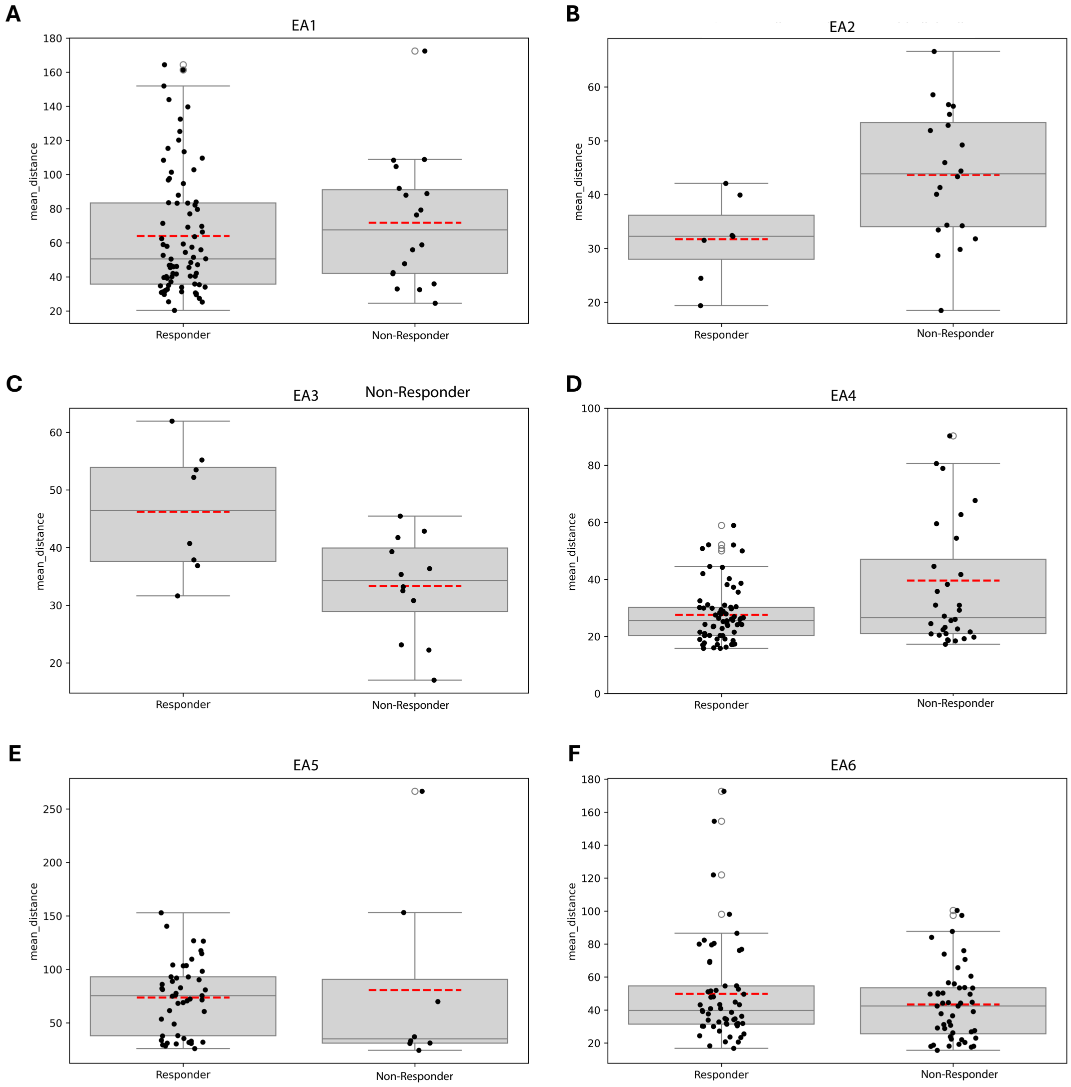


**Fig. S10** Mean Distances Between Immune and Epithelial Cells in Different Epithelial Archetypes: Boxplots comparing the mean distance between epithelial and immune cells within ROIs assigned to Epithelial Archetypes (EA1–6; A–F), for responders and non-responders.

### References

[1] Allen, H.C., Sharma, P.: Histology, Plasma Cells. In: StatPearls. StatPearls Publishing, Treasure Island (FL) (2025). http://www.ncbi.nlm.nih.gov/books/NBK556082/ Accessed 2025-08-27
